# Architecture and function of Bud3 and Bud4-induced septin structures

**DOI:** 10.1101/2025.10.03.680245

**Authors:** Ingrid E. Adriaans, Sandy Ibanes, Aurélie Bertin, Chantal Cazevieille, Joséphine Lai-Kee-Him, Marco Geymonat, Laura Picas, Simonetta Piatti

## Abstract

At the time of cytokinesis, a double septin ring is assembled at the division site in many eukaryotic cells. In budding yeast the double ring is made by two arrays of circumferential septin filaments at the two sides of the bud neck, which are thought to compartmentalize the membrane around the cleavage site. Integrity of the double septin ring requires the anillin Bud4 and its presumed partner Bud3, which associate with septins in mitosis and have separate, yet unknown, roles in stabilizing septin circumferential filaments.

Through *in vitro* reconstitution assays using purified proteins, we show that Bud3 and Bud4 organise septin filaments in distinct ways, while together they cooperate to assemble higher-order septin networks. In agreement with their separate roles in septin organization, Bud3 and Bud4 bind to different septins and require septins to associate with each other, indicating that they modulate septin architecture independently but synergistically. We also provide evidence that Bud3 and Bud4 bind membranes *in vivo* and *in vitro*, consistent with the presence of lipid-binding domains in their primary sequence.

Using *bud3 bud4* double mutants that lack the septin double ring, we show that *in vivo* proteins marking cytokinetic remnants require the double ring to efficiently concentrate at the division site, while other proteins landing at the bud neck at cytokinesis are unaffected. Thus, we propose that a septin double ring may imprint a selective spatial memory for cytokinesis that is transmitted throughout subsequent cell divisions.

## INTRODUCTION

Septins are a family of cytoskeletal GTP-binding proteins that are found in many eukaryotes ^1^. They form nonpolar hetero-hexamers and -octamers containing two molecules of different septins arranged in a palindromic fashion. In turn, these complexes can polymerize into filaments and/or form higher-order structures such as spirals, rings and gauzes ^2,3^. These septin assemblies are often found in close proximity to the cell membrane, where they are thought to act as diffusion barriers and/or scaffolds for recruitment of other proteins ^4^. As a result, they play important roles in many membrane-linked processes such as cytokinesis, neuronal morphogenesis, sperm maturation and cell migration ^5^.

In budding yeast, five septins (Cdc3, Cdc10, Cdc11, Cdc12 and Shs1) are expressed during the mitotic cell cycle and form rod-like nonpolar octamers, with a core Cdc12-Cdc3-Cdc10-Cdc10-Cdc3-Cdc12 hexamer and either Cdc11 or Shs1 at the ends ^6–8^. While Cdc11-capped octamers can polymerise by end-to-end annealing to form long paired filaments, Shs1-capped octamers rather organise into multi-layered rings of staggered septin rods ^6,7,9^. Two meiotic-specific septins, Spr3 and Spr28, replace Cdc12 and Shs1, respectively, during gametogenesis ^10–13^. The mitotic budding yeast septins, except for Shs1, are essential for cytokinesis by forming a rigid collar (also referred to as “septin hourglass”) that recruits most cytokinetic proteins to the division site ^14,15^. The septin collar is also required for maintenance of cell polarity during cell division, possibly by functioning as a diffusion barrier ^16–18^.

The septin hourglass consists of axial septin filaments oriented along the cell division axis, intersected perpendicularly by a few circumferential septin filaments ^19–21^. Septin filaments are proximal to the plasma membrane, but whether they bind directly to the membrane through their polybasic domain and/or membrane interaction is mediated or reinforced by adaptor proteins is unclear^19,22,23^.

During cytokinesis the septin collar is remodelled into a double septin ring (reviewed in ^24^). The double septin ring is made exclusively by two arrays of circumferential filaments, suggesting that during remodelling axial septin filaments may get selectively depolymerized or displaced and perhaps reassembled into new circumferential filaments ^20,25,26^. Importantly, this septin rearrangement is strictly necessary for cell division, possibly by allowing the cytokinetic machinery that is sandwiched between the two split septin rings to make contact with the plasma membrane and drive cleavage furrow ingression and septum formation ^27,28^. Additionally, the double septin ring has been proposed to create a cortical compartment that constrains cytokinetic proteins at the division site ^16,17^.

Formation and maintenance of the double septin ring require the anillin homologue Bud4 and the Rho-GEF Bud3 ^29–32^. However, the exact molecular mechanism underlying septin remodelling and the assembly of a double septin ring is unknown.

Bud3 and Bud4 had been originally characterised for their role in defining a cortical landmark for the axial budding pattern of haploid yeast cells ^33^. While wild type haploid cells bud in an axial manner (i.e. with the new bud emerging next to the old bud site), in the absence of Bud3 or Bud4 they bud in a bipolar fashion (i.e. with the new bud appearing on the opposite side of the mother cell relative to the previous bud site), similar to diploid cells ^34^.

Bud3 and Bud4 interact with each other and with septins ^31,32,35–37^. In addition, both proteins have membrane-binding motifs, i.e. a pleckstrin-homology (PH) domain at the C-terminus of Bud4 and an amphipathic helix located in the middle of Bud3 ^29,31,38,39^. Overexpression of either protein leads to the formation of supernumerary septin rings and arcs (Bud4) or spirals (Bud3) at the cell membrane ^31,39,40^. In addition, the amphipathic helix of Bud3 is required to induce extra septin spirals upon Bud3 overexpression ^39^.

Bud3 and Bud4 are first recruited to the mother side of the septin collar during G2/M and, shortly prior to septin remodelling into a double ring, they are also found on the daughter side of the septin collar. After septin remodelling, they remain associated with the double septin ring until the next G1 phase ^29,37,41–43^.

Mutants lacking *BUD4* do not form a double septin ring but disassemble the septin hourglass at cytokinesis, especially on the mother side of the bud neck. Deletion of *BUD3* causes instead the fragmentation of the septin collar at cytokinesis, while combined deletion of *BUD3* and *BUD4* virtually eliminates all septin structures at the bud neck at cytokinesis. In addition, ultrastructural data showed that Bud3 and Bud4 have overlapping, yet distinct roles in ensuring the integrity of the “transitional” hourglass (i.e. just before septin remodelling) and together provide positional cues for the assembly of the double septin ring ^29^. Based on all these observations, it is reasonable to assume that Bud3 and Bud4 stabilize the circumferential septin filaments at the plasma membrane during cytokinesis ^44^. However, the mechanistic details of how this is achieved and whether the membrane-binding domains of Bud3 and Bud4 are involved remain to be investigated.

Using *in vitro* reconstitution assays based on recombinant proteins either in solution or on supported lipid bilayers, we show how Bud3 and Bud4 promote septin organization and interaction with the cell membrane. In addition, to gain further insights into the proposed role of the septin double ring as a membrane diffusion barrier, we took advantage of the phenotype of *bud311 bud411* double mutants, i.e. complete absence of a double septin ring, to assess which proteins, among those appearing at the division site around the time of cytokinesis, require the septin double ring to accumulate at the bud neck.

## RESULTS

### Bud3 and Bud4 cooperatively assemble higher-order septin structures *in vitro*

To study how Bud3 and Bud4 directly impact septin organization, we set up *in vitro* reconstitution assays. To facilitate protein purification, we reasoned that the protein domain of Bud4 involved in septin binding and organisation would lie within its region of homology with its fission ortholog *S.p.* Mid2, which has similar septin-related properties to Bud4 ^45,46^. Thus, we expressed and purified from bacterial cells a short version of Bud4 (aa 623-1447, henceforth called Bud411N), as well as Cdc11-Cdc12-Cdc10-Cdc10-Cdc3-Cdc12-Cdc11 septin octamers ^47,48^ (Fig. 1A, S1A). To allow polymerization, septins were shortly incubated in low salt buffer (30 mM NaCl) in the absence or presence of Bud411N. Septin filaments were visualized by Transmission Electron Microscopy (TEM) after negative staining (Fig. 1B). Septin octamers without Bud4 formed paired septin filaments, as previously reported ^6^. In the presence of Bud411N, septin “ladders” with periodic lateral bridges between paired filaments were apparent under the same conditions (Fig. 1B). These septin ladders were reminiscent of previously described railroad-like septin structures that were formed in the presence of Bni5, Gic1 or Shs1 ^7,49,50^. Rarely, we observed them also with Cdc11-capped octamers in the absence of any other protein (data not shown), suggesting that the lateral crosslinks may be generated by one of the septins, aided by different septin-binding proteins. The average distance between each step of the septin ladder structure was approximately 33 nm (Fig. 1B-C), which corresponds to the length of one octameric rod ^6^. Thus, Bud4 may stimulate the formation of septin ladders. Since the septin-binding region of Bud4 (aa 623-774) was previously shown to be in close proximity to Cdc11, Shs1 and Cdc3 *in vivo* ^51^, in our *in vitro* assays Bud4 must promote lateral bonds between paired septin filaments by modulating the conformation of Cdc11 or Cdc3 (Shs1 is absent from Cdc11-capped octamers). However, given that the step size is constant and has the length of one octamer, it is likely that Bud4 binding to Cdc11, rather than Cdc3, may induce septin bridging.

**Figure 1.**
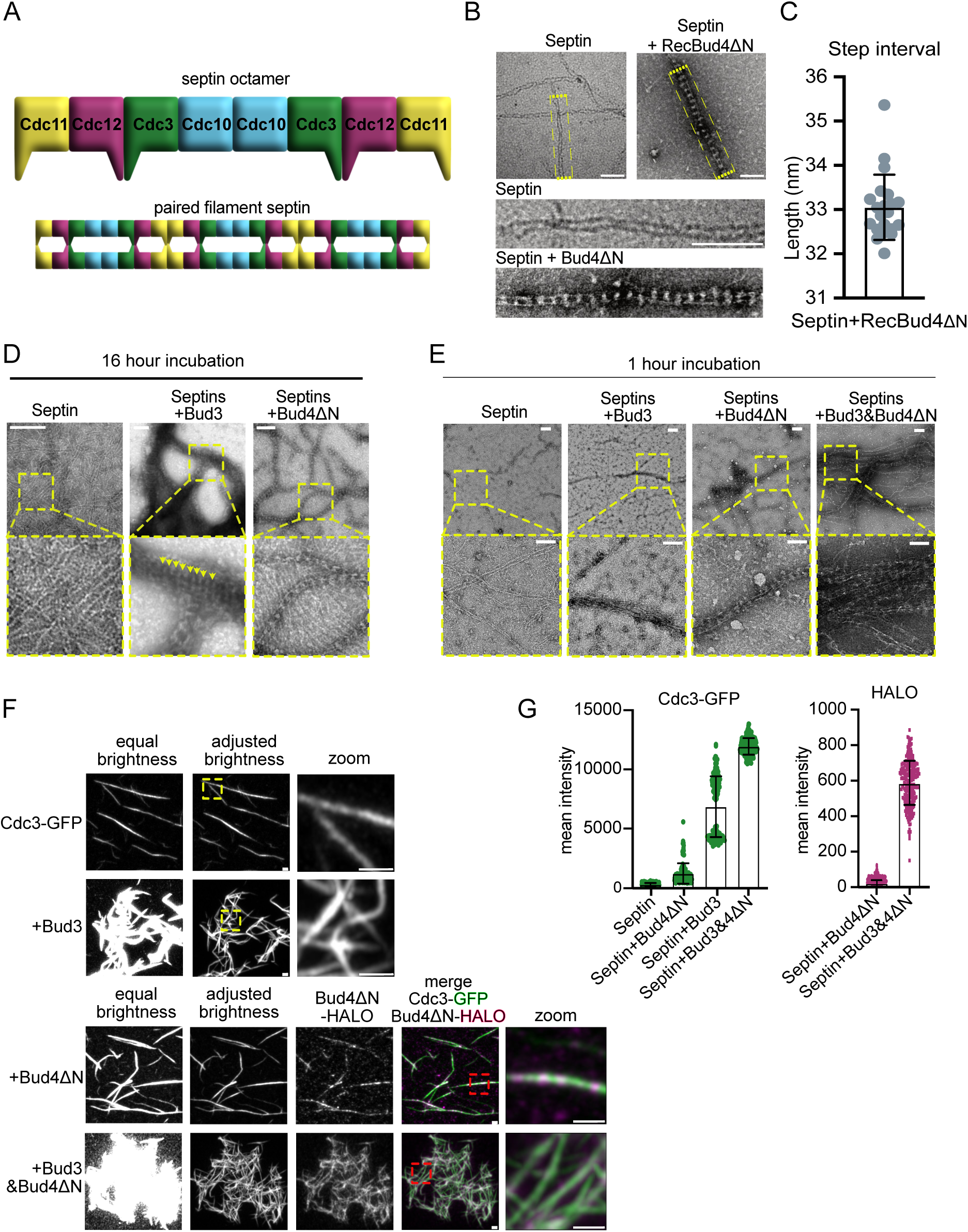
Bud3 and Bud4 organize septin filaments into higher order structures. **A:** Schematic representation of a budding yeast Cdc11-capped septin octamer and a paired septin filament. **B-C:** 90 nM septin octamers were polymerized in solution by lowering the salt concentration (50 nM NaCl) and incubating for 30 minutes on ice in the absence or in the presence of equimolar amounts of recombinant Bud4ΔN. Representative TEM images, including magnified insets (yellow boxes), are shown in B. Scale bars: 100 nm. Note the periodicity of septin lateral bridges in the presence of Bud4ΔN protein is added, whose intervals are shown in C (n=21). **D-E:** 180 nM septin octamers were polymerized in solution by lowering the salt concentration (30 nM NaCl) in the absence or in the presence of 90 nM of Bud3 or Bud4ΔN, or 45 nM of Bud3 and Bud4 purified from *S. cerevisiae*. TEM images were taken after 16 hours or 1 hour of incubation at 4°C. Yellow boxes indicate magnified insets that are shown in the bottom row. Arrowheads indicate periodic striations in septin bundles. Scale bars: 200 nm. **F-G:** 40 nM septin octamers were incubated with 20 nM of Bud3 and/or 20 nM of Bud4ΔN-HALO in low salt buffer (30 mM NaCl) for 1 hour at 4°C. Zoom areas are indicated with yellow and red boxes. Scale bars are 1 µm. Fluorescence intensities of Cdc3-GFP (n=125, 143, 179 and 236, respectively) and Bud4ΔN-HALO (n=274 and 261, respectively) for the indicated conditions have been plotted in G. Mean values and standard deviations are shown.

Bud3 is a large protein of 185 kDa, making its purification from *E. coli* very challenging. In addition, the quality of our purified Bud411N from bacterial cells was not optimal due to the presence of degradation products (data not shown). Thus, we purified MBP-tagged full-length Bud3, as well as Bud411N, from budding yeast (Fig. S1B-C). After sixteen hours of incubation in low salt buffer, septin octamers alone had formed long paired filaments, as expected, while the presence of Bud4-11N purified from yeast induced the formation of septin ladders similar to the ones we observed with recombinant Bud411N (Fig. 1D). Conversely, Bud3 formed large and dense septin structures made out of bundled filaments. In some areas the Bud3-induced septin bundles showed clear parallel periodic striations (Fig. 1D, yellow arrowheads). We were unable to image septin structures formed by Bud3 and Bud4 together under the same conditions, as the EM grids would get destroyed before or during imaging by massive amounts of highly dense septin structures. Therefore, we reduced the concentration of Bud3 and Bud4 and decreased the incubation time to one hour. Septins alone or septins in the presence of Bud411N formed similar structures as before, although in the latter condition septin ‘ladders’ were fewer and more dispersed on the grid than after 16 hours of incubation (Fig. 1E). In the presence of Bud3, relatively thin bundles of septin filaments were apparent (Fig. 1E), suggesting that prominent bundling depends on high protein concentrations and/or long incubation times. However, when Bud3 and Bud411N were added together to septins, large bundled septin ‘ladders’ readily formed (Fig. 1E), indicating that Bud3 and Bud4 cooperate to organise septins in higher-order structures.

In order to quantify septin assemblies induced by Bud3 and/or Bud4, we imaged eGFP-tagged septin octamers polymerised *in vitro* in low salt buffer for one hour in the absence or presence of Bud3 and/or Alexa-Fluor®660-labeled Bud411N-HALO by fluorescence microscopy (Fig. 1F). Using this set-up, septin filaments alone looked similar to the ones previously reported on PEG-treated glass ^8^. Addition of Bud3 increased drastically the fluorescence intensity of septin structures and generated large septin networks, probably due to septin filament bundling that we observed by TEM. Unfortunately, we did not succeed to fluorescently tag Bud3 to visualize its colocalization with septins, as Bud3 would no longer induce septin networks after HALO tagging (data not shown). Conversely, functional binding of Bud411N-HALO to the red fluorescent Alexa-Fluor®660 ligand enabled us to quantify the fluorescence of septin assemblies, as well as the Bud4 signal associated to septin structures (Fig. 1F, G). Addition of fluorescent Bud411N-HALO to septins caused a modest increase in septin fluorescence intensity and no clear change in morphology (Fig. 1F). This is again consistent with our TEM results, as the septin ladders would be indistinguishable from paired septin filaments at this magnification. Conversely, co-addition of Bud3 and fluorescent Bud411N-HALO to polymerizing septin octamers resulted in the formation of large septin networks with an architecture similar to the one observed in the presence of Bud3 alone, but more densely packed (Fig. 1F). Additionally, these meshes displayed greatly enhanced fluorescence intensity of both Cdc3-GFP and Alexa-Fluor®660-labeled Bud411N-HALO (Fig. 1G), confirming that Bud3 and Bud4 cooperate in a synergistic manner to form septin supra-molecular assemblies.

### Bud3 interacts with Cdc10 *in vivo*, and septin polymerization mediates Bud3–Bud4 binding

In order to learn more about how Bud3 and Bud4 work together to form higher-order septin structures, we used a tripartite GFP system ^52^. Briefly, in the tripartite GFP system two potentially interacting partners are tagged, respectively, at the C-terminus with the β11 strand of GFP (21 residues) and at the N-terminus with the β10 strand of GFP (20 residues). In case of protein-protein interaction, reconstitution of GFP is possible thanks to the capture of the non-fluorescent GFP β1-9 barrel co-expressed from a plasmid by the galactose-inducible *GAL1* promoter ^51^. This strategy was used to show that Bud4 contacts physically the septins Cdc3, Cdc11 and Shs1 in yeast cells ^51^. In order to study the interaction of Bud3 with septins in yeast cells, we tagged full-length Bud3 at its C-terminus with GFP β11, while the β10 GFP strand was fused to the N-terminus of each mitotic septin ^51^ (Fig. 2A). Upon expression of GFP β1-9, β10-Cdc10 was the septin that in combination with Bud3-β11 gave the strongest fluorescent signal at septins, marked by Cdc11-mCherry (Fig. 2B). This indicates that Bud3 is in close proximity to or interacting with Cdc10 and not with the other septin subunits. This result is in agreement with the previous finding that Bud3 pulls down preferentially Cdc10 from yeast extracts compared to the other septins ^39^. The tripartite GFP interaction could not be detected in unbudded or small budded cells (i.e. in G1 or S phase), consistent with the mitotic expression of Bud3 ^29,43,53^, while it was strong at the bud neck of large budded cells, both before and after septin remodelling (Fig. 2B-D). In addition, it decorated the opposed edges of the septin collar in mitosis and the split septin rings, in line with the previously reported localization of Bud3 ^29,43,53^. Thus, Bud3 interacts with Cdc10 from S/G2 until the completion of cytokinesis.

**Figure 2.**
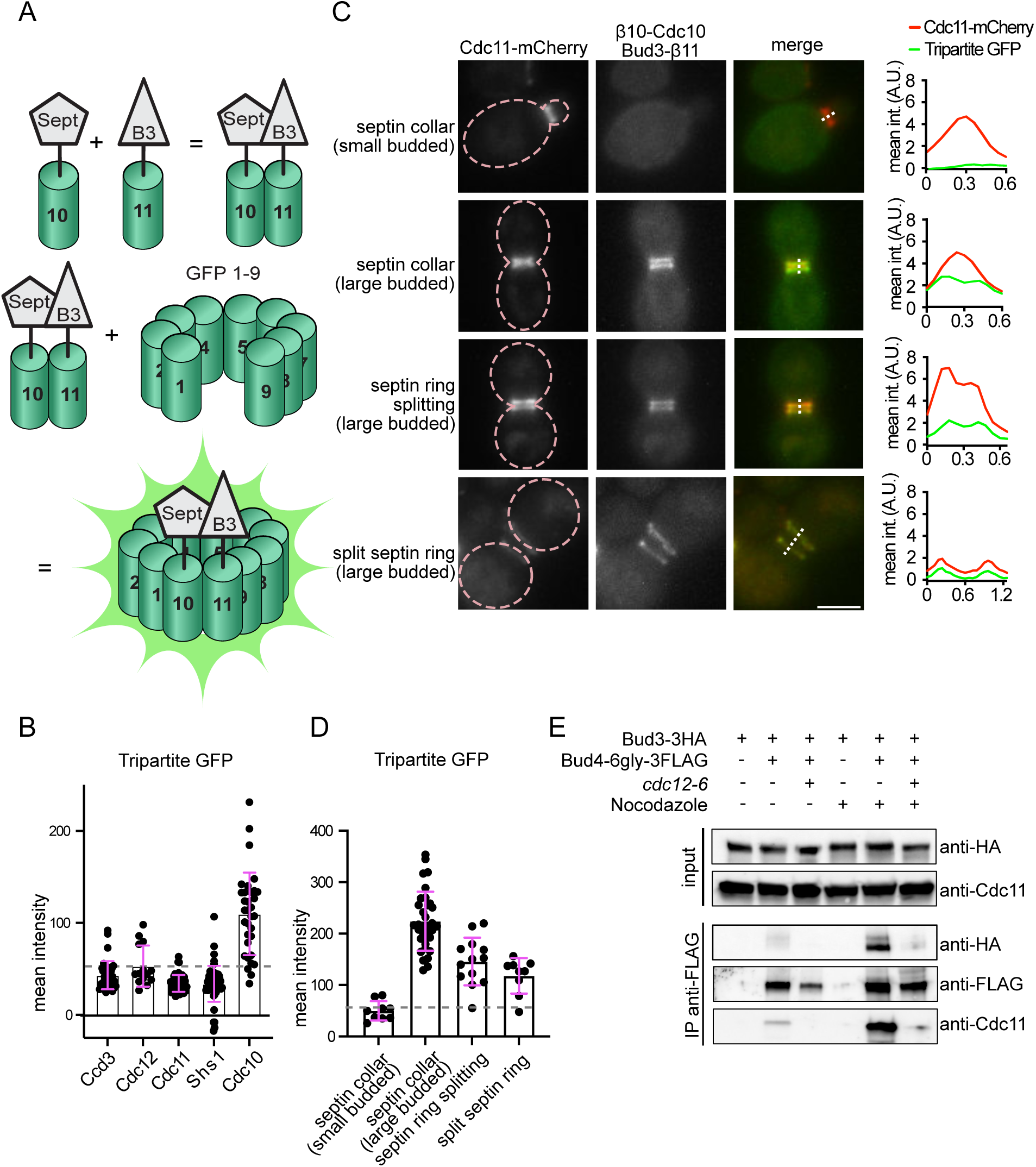
Bud3 interacts directly with the septin Cdc10, while its interaction with Bud4 depends on septins. **A:** Schematic view of the tripartite GFP system. B3: Bud3 and Sept: Septin. The GFP β10 and β11 strands are fused to the N-terminus of Cdc10 and the C-terminus of Bud3, respectively. Upon expression of the GFP β1-9 barrel from the galactose-inducible *GAL1* promoter, the close proximity of β10 and β11, together with β1-9, will reconstitute fluorescent GFP. **B:** Quantification of fluorescence intensities of the tripartite GFP signal at the septin collar when β10 is fused to different septins. Measurements were done in large budded cells with a septin collar, i.e. at cell cycle stages when Bud3 is maximally expressed, before septin remodelling. The background GFP signal was measured away from the septin collar but inside the cell and subtracted from the values at the septin collar. Some signals at the bud neck were higher than the background but present in all cells. Thus, they were considered unspecific. A dotted line indicates the cut-off value that was set at 50. Mean values and standard deviations are shown in pink. n = 38, 14, 50, 54 and 33, respectively. **C:** Representative images of yeast cells expressing Cdc11-mCherry, β10-Cdc10, Bud3-β11 and GFP β1-9 at different stages of the cell cycle. Scale bar: 2 µm. GFP and mCherry fluorescence was measured along the dotted line indicated in the merge images and plotted in the graphs shown on the right. Mean int. = mean intensity and A.U.=arbitrary units. **D:** Quantification of fluorescence intensities of the tripartite GFP signal due to Bud3-Cdc10 interaction at the bud neck and at different cell cycle stages. Background values (inside the cell but away from the bud neck) were subtracted from fluorescence intensities at the bud neck. The dotted line indicates the cut-off value that was set at 50. Mean values and standard deviations are shown in pink. n = 9, 33, 13 and 9, respectively. **E:** Asynchronously growing or nocodazole-arrested wild type and *cdc12-6* cells expressing the indicated tagged proteins were shifted to 37°C for 30 minutes to depolymerise septins in *cdc12-6* cells. Protein extracts were subject to immunoprecipitation with anti-FLAG antibody. Total extracts (inputs) and immunoprecipitates (IP) were loaded on SDS page for western blot detection of Bud3-3HA and Cdc11.

Bud3 and Bud4 are thought to form a complex ^31^, although we show here that Bud3 interacts with a different septin relative to Bud4 and that Bud3 and Bud4 act synergistically on septin organisation. Therefore, we wondered if the interaction between Bud3 and Bud4 may be mediated by septins. To this end, we compared the amount of HA-tagged Bud3 (Bud3-3HA) co-immunoprecipitated with Flag-tagged Bud4 (Bud4-6Gly-3Flag) in wild type and *cdc12-6* cells, which are known to quickly depolymerize septins at the bud neck upon a short incubation at high temperatures (30°-37°C) ^16,17^. Low amounts of Bud3-3HA could be co-immunoprecipitated with Bud4-6Gly-3Flag in logarithmically growing wild type cells, indicative of Bud3 binding to Bud4. Interaction was dramatically increased in cells enriched in mitosis by nocodazole treatment (Fig. 2E), in agreement with the mitotic expression of both proteins. In both cycling and nocodazole-arrested cells, septin depolymerisation by the *cdc12-6* allele disrupted the interaction between Bud3-3HA and Bud4-6Gly-3Flag (Fig. 2E). Concomitantly, also interaction between Bud4-6Gly-3Flag and the septin Cdc11 was severely impaired, suggesting that Bud4 recognizes septin filaments, rather than octamers. Thus, we conclude that interaction between Bud3 and Bud4 depends on the integrity of septins filaments at the bud neck. However, we do not rule out that Bud3 and Bud4 also interact with each other upon their association with septins.

### Bud3 and Bud4 together help large septin assemblies to bind the membrane

Septin structures are often found in close proximity to the plasma membrane ^4^ and septins, Bud3 and Bud4 have membrane-binding domains. Thus, we asked whether and how Bud3 and Bud4 would organize septins on membranes. We used an *in vitro* reconstitution assay using Supported Lipid Bilayers (SLBs) and purified GFP-tagged Cdc11-capped recombinant septin octamers ^47,48^, as well as MBP-tagged Bud3 and Bud411N purified from yeast, as mentioned above. We used a lipid composition of 87 mol% phosphatidylcholine (PC), 2.8 mol% PI(4,5)P_2_, 10 mol% phosphatidylinositol (PI) and 0.2 mol% fluorescent PI(4,5)P_2_ (TF-TMR-PI(4,5)P_2_) to visualize the membrane. Upon a short (∼5 min) incubation in low salt buffer, septins were injected onto the SLB, leading to the appearance of sub-micrometric septin filaments on the SLB within minutes (Fig. 3A). Sparse filaments adhered to the SLB along their entire length, while most of them bound the SLB only with one end while remaining floppy in the bulk (Movie S1). Co-addition of Bud3 or Bud411N interfered with septin binding to the SLB (Fig. 3A, with Bud3 leading to formation of large septin structures in solution above the membrane (similar structures as in Fig. 1F), suggesting that either septins have higher affinity for Bud3 and Bud4 than for lipids, or Bud3 and Bud4 compete with septins for membrane binding. In contrast, upon co-addition of Bud3 and Bud411N longer septin bundles appeared on SLBs than with septins alone (Fig. 3A), confirming that Bud3 and Bud4 cooperate in the organisation of septin higher order structures and suggesting that they may help membrane binding of septin filaments along their length. Quantification of filaments on the SLB and in the maximum 3 µm^3^ volume above the membrane with SOAX revealed that their median length increased significantly in the presence of Bud3 and Bud4ΔN (0.64±0.42 µm, n=8646 for septins alone; 1.21±0.92 µm, n=5610 for septins with Bud3 and Bud4ΔN) (Fig. 3B and S2A), while their curvature remained similar (0.26±0.23 µm^−1^, n=85 for septins alone; 0.23±0.21 µm^−1^, n=1138 for septins with Bud3 and Bud4ΔN) (Fig. 3C). We reproducibly noted that in the presence of Bud3 and/or Bud4 the fluorescent signal of PI(4,5)P_2_ at the SLB decreased significantly, but at the moment we do not know whether this is due to stripping, clustering or quenching of PI(4,5)P_2_ (Fig. S2B).

**Figure 3.**
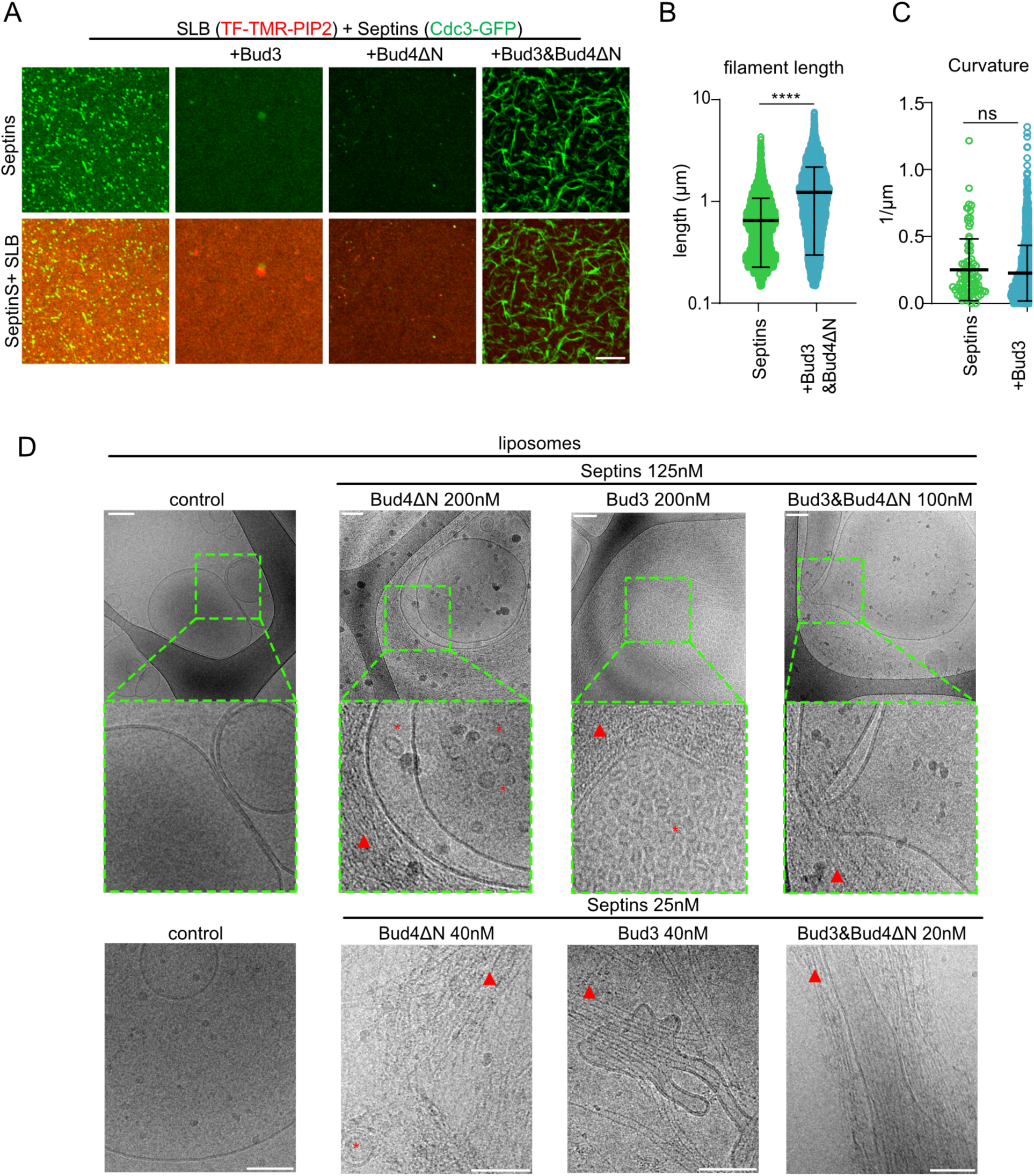
Bud3 and Bud4 together favours membrane binding of septin filaments. **A:** Representative images of SLBs incubated with 40 nM GFP-tagged septin octamers with or without 20 nM of Bud3 and/or 20 nM of Bud4ΔN-HALO in low salt buffer (30 mM NaCl). SLBs are visualized with TF-TMR-PIP2 and septins by Cdc3-GFP. Scale bar: 5 µm. **B-C:** Distribution of lengths (B) and curvatures (C) of septin structures calculated by the SOAX software (B n=8646 for septins alone and n=5610 for septins with Bud3 and Bud4ΔN, C n=85 for septins alone and n=1138 for septins with Bud3 and Bud4ΔN). Only the conditions where septin filaments were present on the SLB were analysed. **D:** The indicated concentrations of septins were added to liposomes in the absence or presence of the indicated amounts of Bud4ΔN, Bud3 or both. Septin filaments and liposomes were visualized by CryoEM. Green boxes indicate magnified insets shown in the row below. Asterisks indicate small lipid vesicles, red arrowheads indicate septin bundles. Scalebar: 100 nm.

To look more closely at Bud3 and Bud4-induced septin assemblies on lipids, we visualized septin assemblies on lipid monolayers by TEM after negative staining ^54^. On lipid monolayers septin octamers formed paired and striated filaments (Fig. S2C) similar to the ones was previously described ^54^. Addition of Bud3 or Bud411N regardless of the presence of septins caused prominent monolayer deformations that are indicative of their interaction with lipids, but no septin filaments or bundles could be observed (Fig. S2C).

Next, we visualised septin assemblies on liposomes by CryoEM. Using different protein concentrations of septins together with Bud3, Bud411N, or the two proteins together, we observed tightly packed septin bundles made of laterally ordered filaments (Fig. 3D). Septin assemblies induced by Bud3 and/or Bud411N that were in solution, i.e. away from liposomes, showed bundles made of evenly spaced septin filaments (Fig. S2D). Septin-structures induced by both Bud3 and Bud4 seemed more densely packed, forming mesh-like structures (Fig. S2D).

Interestingly, at high concentrations Bud3 and Bud411N also caused the formation of many small lipid vesicles, presumably as a result of their ability to bind and deform membranes (Fig. 3D, red asterisks).

Thus, we conclude that in spite of having lipid-binding domains, Bud3 or Bud4 separately induce septin higher-order structures mainly in solution, away from lipids, while the two proteins together favour membrane interaction of septin filaments. It remains however possible that the lipid composition and/or lack of curvature of our SLBs or lipid monolayers is not suited to recapitulate the membrane interaction of septin structures induced by Bud3 or Bud4.

### Bud3 and Bud4 remodel membranes partly via their PH and AHX domain

We noticed that when we added Bud3 or Bud411N to SLBs we could observe membrane deformations, such as vesicles and tubules growing out of the SLB plane. These were difficult to image because they were very dynamic and growing in the z-plane (data not shown). To better visualize them, we used silica beads coated with a lipid bilayer of the same composition as in the experiments described above that enabled us to look at the deformations in one z-plane. Using this set-up we could nicely visualize the formation of tubules induced by Bud3 or Bud411N (Fig. 4A), suggesting that Bud3 and Bud4 interact with and deform the membrane.

**Figure 4.**
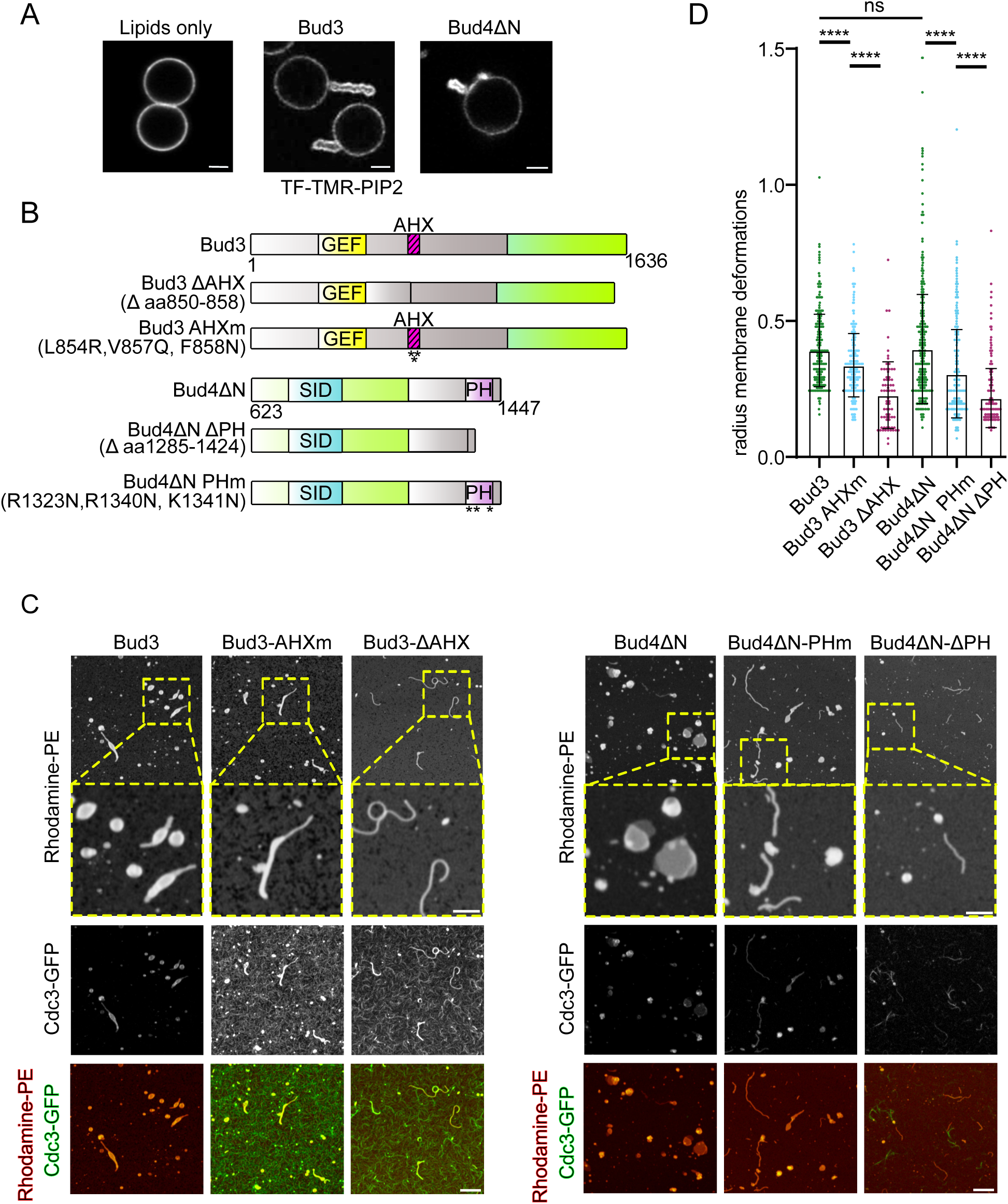
Bud3 and Bud4 induce membrane tubulation. **A:** Representative images of SLBs on 5 µm diameter silica beads visualised by TopFluor®-TMR-PI(4,5)P2 either in the absence or in the presence of 40 nM Bud3 or Bud4ΔN. Scale bar: 2 µm. **B:** Schematic representation of the different Bud3 and Bud4ΔN mutant proteins used in C and D. GEF=Guanine nucleotide Exchange Factor, AHX=Amphipathic Helix, SID=Septin Interacting Domain, PH=Pleckstrin Homology domain. **C:** Representative images of membrane deformations of an SLB with a composition of 95.8 mol % DOPC, 4 mole % DGS-NTA(Ni) and 0.2 mol % Rhodamine-PE induced by 40 nM/each Bud3 and Bud4ΔN visualized by Rhodamine-PE and Cdc3-GFP. Scale bar: 2 µm. **D:** Quantification of the radius of membrane deformations from the experiment in C. n= 284, 303, 70, 554, 472 and 292.

The PH domain of Bud4 was shown to bind to some classes of phosphoinositides ^38^, while Bud3 bears an amphipathic helix in its middle region that together with an adjacent stretch of basic residues can interact with the plasma membrane ^39^. To investigate if and how these protein domains impact the interaction of Bud3 and Bud411N with membranes *in vitro*, we either deleted them or introduced specific mutations ^38,55^ (Fig. 4B). In the case of Bud3, we mutated the conserved hydrophobic residues of the amphipathic helix ^39^, while for Bud411N we mutated the conserved hydrophobic residues of the PH domain that were reported to convey membrane interaction of phospholipase C-delta 1 ^56^. Since preliminary experiments indicated that membrane protrusions induced by Bud3 (Movie S2) or Bud411N (Movie S3) were highly mobile and flexible in 3D and that septins would bind to these tubules, in order to quantify the extent of membrane deformation, we immobilized membrane protrusions using nickel salt-containing SLBs and 6His-GFP-tagged septin octamers. The 6His-tagged septin octamers would bind to membrane protrusions induced by Bud3 or Bud411N and stick to the nickel salts in the SLB. Since MgCl_2_ is necessary for GTP binding and septin bundling ^8^, we omitted it from these assays. Both Bud3 and Bud411N (40 nM each) prompted the formation of membrane vesicles and tubules (Fig. 4C). Mutation or deletion of the membrane binding domains of Bud3 and Bud4 did not abrogate membrane deformations, suggesting the possible presence of other membrane binding domains in the two proteins. However, they did alter the shape, mainly the curvature (radius), of membrane protrusions, which became more tubular and thinner (Fig. 4C-D). Indeed, in the presence of intact Bud3 or Bud411N the curvature of membrane deformations was shallower (i.e. higher radius) and significantly increased with the Bud3 AHX or Bud4 PH mutant proteins. Protrusions and tubules in SLBs can be induced by the expansion of the bilayer and likely stem from Bud3 and Bud4 membrane binding. However, the curvature of membrane deformations may be specified by the PH domain of Bud4 and the AHX domain of Bud3. Interestingly, we observed a clear increase in septin filaments on the SLB in the presence of either Bud3 mutant protein (Fig. 4C), suggesting that the AXH may affect the affinity of Bud3 for septins.

### The PH domain of Bud4 is essential for septin remodelling *in vivo*

Deletion of *BUD4* disrupts the formation of the septin double ring in yeast cells, while deletion of *BUD3* has a milder effect ^29,31,32,57^. In order to delete the PH domain of Bud4 (aa 1286-1447) and determine its impact on septin remodelling at cytokinesis, we inserted the eGFP after aa 1285 at the *BUD4* endogenous locus, thus creating a deletion of the last 162 amino acids encompassing the PH domain. In cells expressing full-length Bud4-eGFP, the septin collar was split in two between mother and daughter cell, as expected. Additionally, Bud4-eGFP was recruited to the septin collar in medium/large budded cells (i.e. in mitosis) consistent with its transcription pattern ^58^, and colocalized with the septin double ring during and after cytokinesis, as previously reported ^29,31,32,42^ (Fig. 5A). Conversely, when the PH domain was removed, septins got quickly dismantled at cytokinesis, most notably on the mother side of the bud neck, and the septin double ring never formed (Fig. 5B). This phenotype is highly reminiscent of that of a mutant previously described bearing a deletion that encompasses the entire C-terminus (anillin homology and PH domains, aa 1067-1447) ^29^, indicating that the PH domain is a major determinant in septin double ring integrity. Contextually, Bud4-11PH-eGFP still co-localized with septins (Fig. 5B), in line with the conclusion that the septin-interacting domain of Bud4 lies in the middle of the protein, probably aided by the C-terminal anillin homology and/or PH domain ^29,31,32^.

**Figure 5.**
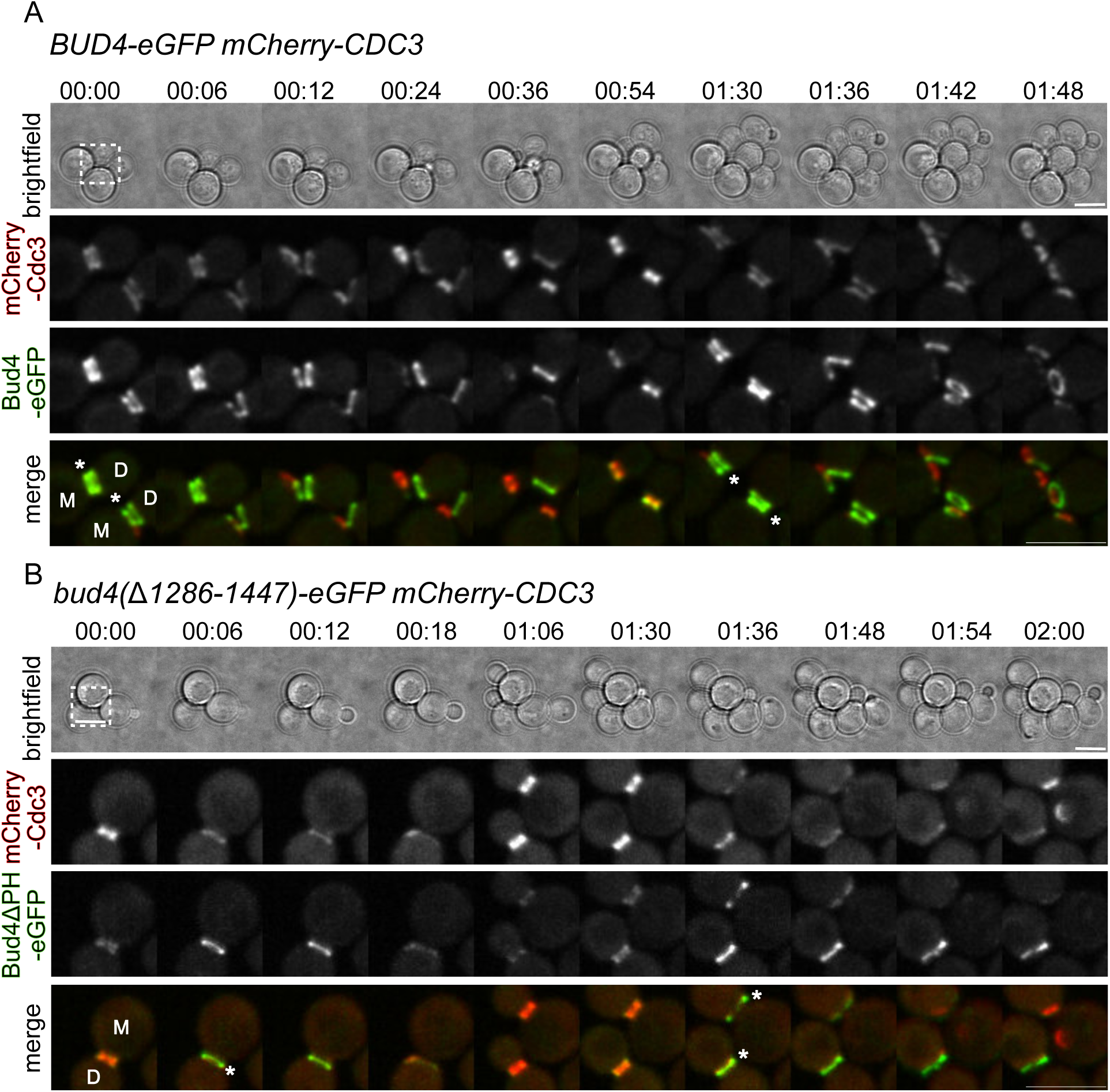
The PH domain of Bud4 is necessary for formation of the double septin ring. **A-B:** Cells with the indicated genotypes were imaged at 30°C every 6 minutes. Time is indicated in hours:minutes. Asterisks indicate septin reorganization at cytokinesis. M: mother cell, D: daughter cell. Scale bar: 2 µm.

Bud3 and Bud4 tagged with mNeonGreen and 4XGFP, respectively, co-localized with the septin collar in mitosis, and kept rising at the bud neck even after septin remodelling (Fig. S3A), suggesting that their membrane-binding domains may mediate bud neck localisation after septin depolymerisation at cytokinesis. Consistent with and extending previous findings ^29,30,37^, deletion of *BUD4* (*bud4Δ*) or the *bud4* frameshift mutation (G2459fs) found in the W303 background that truncates the C-terminus of Bud4 after aa 870, led to a severe decrease of Bud3 localization at the bud neck (Fig. S3B). Additionally, localization of Bud4(G2459fs)-4GFP at the bud neck was drastically compromised (Fig. S3C), in agreement with the idea that the minimal septin-interacting domain alone (aa 623-774) is not sufficient for efficient Bud4 interaction with septins ^29,31,32,51^, which needs also the anillin homology and/or PH domain.

Deletion of *BUD3* did not affect the localization of Bud4 prior to cytokinesis, but resulted in aberrant septin remodelling, as described before ^29^ (Fig. S3C). Thus, Bud4 localizes to septins mainly independently of Bud3, while Bud3 localization prominently requires Bud4. However, the already impaired localization of Bud4(G2459fs)-4GFP at the bud neck is completely abolished in the absence of Bud3 (Fig. S3C), suggesting that recruitment of Bud3 and Bud4 to the bud neck and septin structures is tightly linked and, to a certain extent, interdependent.

Since Bud3 and Bud4 are recruited to the bud neck in mitosis, we asked if they could be involved in stabilizing the septin collar already during mitosis. To this end, we performed Fluorescent Recovery After Photobleaching (FRAP) in cells expressing mCherry-Cdc3 and arrested in metaphase by *MPS1* overexpression ^59^. Upon bleaching half of the septin collar in wild type cells, very little fluorescence recovery was observed and the little recovery that occurred was in the form of a double septin ring (Fig. S4), in line with previous data ^20,60,61^. This suggest that circumferential septin filaments are already present in the septin hourglass in metaphase and that the turnover of septins in the septin collar only concerns circumferential, as opposed to axial septin, filaments. Thus, circumferential septin filaments appear to be more dynamic than axial filaments, thus explaining the dynamic state of the septin double ring that is exclusively made by circumferential filaments ^20,60,61^. To our surprise, under these conditions, fluorescence recovery in double septin rings prior to cytokinesis was not dependent on Bud3 and Bud4 (Figure S4), suggesting that in metaphase Bud3 and Bud4 have no role in organizing the circumferential filaments in the septin hourglass.

### At cytokinesis Bud3 and Bud4 localise to the bud neck partly independently of septins

Bud3 and Bud4 recruitment to the bud neck was previously shown to be septin-dependent ^42,53^. However, the persistence of Bud3 and Bud4 at the bud neck after septin disassembly at cytokinesis ^29^ (and Fig. S3A) suggests that these proteins may also bind to the bud neck membrane independently of septins. To address this possibility, we assessed Bud3 and Bud4 localisation in a *cdc11* mutant that lacks the N-terminal polybasic domain (*cdc11-ΔPB*) and was recently shown to undergo complete septin disassembly instead of remodelling at cytokinesis ^62^. Quantification of fluorescent signals at the bud neck in twenty cells undergoing cell division showed that in wild type cells, the septin mCherry-Cdc3 underwent a stereotypical and sudden drop in protein levels at cytokinesis (by 30-40%) that accompanies septin rearrangement into a double ring ^27,29,30^. In contrast, septins quickly vanished in the *cdc11-ΔPB* mutant at the time of septin remodelling, reaching fluorescence intensities close to the cytoplasmic background (Fig. 6). Despite septin disappearance, Bud3-mNeonGreen and Bud4-4GFP were retained at the bud neck for a longer time, though to slightly lower levels in the *cdc11-ΔPB* mutant than in wild type cells (Fig. 6). We therefore conclude that Bud3 and Bud4 bud neck localisation is partly independent of septins, at least during cytokinesis, and may involve their membrane-binding properties. We do not exclude, however, that Bud3 and Bud4 recruitment by septins in mitosis could enhance their local concentration to allow their membrane binding at the bud neck.

**Figure 6.**
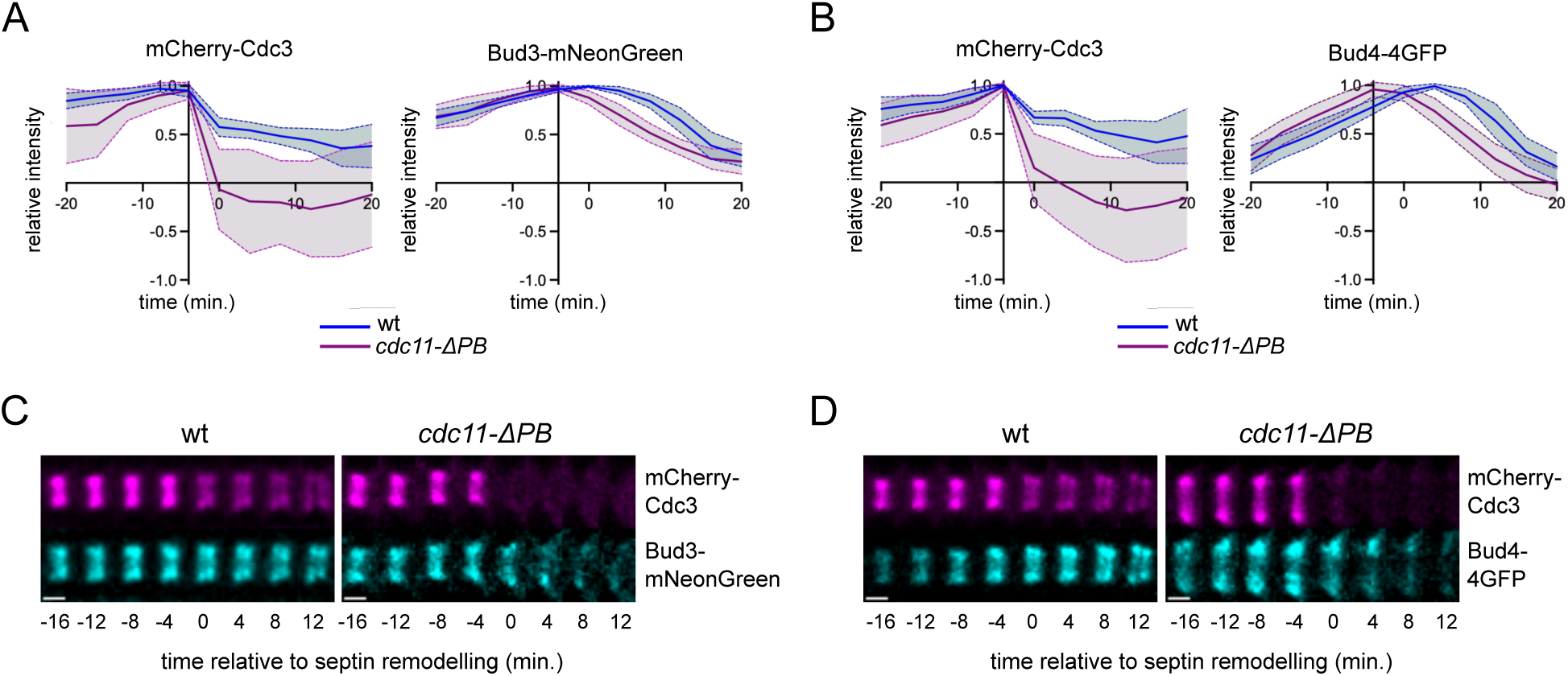
Bud3 and Bud4 are retained at the bud neck after septin depolymerisation. Wild type or *cdc11-!1PB* cells expressing mCherry-Cdc3 and either Bud3-mNeonGreen (A, C) or Bud4-4GFP (B, D) were filmed at 30°C every 4 minutes to quantify mean fluorescence intensities over time as detailed in Materials and Methods. Values were averaged and plotted. Shaded curves: standard deviations. Representative frames of fluorescent signals at the bud neck are shown in C-D. Scale bar: 1 µm.

### A Bud3/Bud4-dependent septin double ring supports protein retention at the division site

The complete disappearance of septins in *bud311 bud411* double mutants at the time of cytokinesis ^29^ allowed us to rigorously assess the involvement of the septin double ring in a membrane diffusion barrier at the division site. We set out to investigate which proteins, among those that are recruited to the bud neck around the time of septin remodelling, depend on the putative barrier function of the septin double ring for their localization. To this end, we analysed by live cell imaging the distribution of three members of the polarisome (Kel1, Spa2 and Bni1), one exocyst subunit (Sec3) and three polarity proteins that are involved in the control of bud site selection (Aim44, Nba1 and Nis1) ^17,63–70^. These selected proteins were tagged with GFP and expressed at endogenous levels alongside the septin Cdc3 tagged with mCherry. After filming wild type and *bud311 bud411* double mutants, we measured the fluorescence intensity of each protein before and after septin remodelling. While in wild type cells during septin remodelling mCherry-Cdc3 protein levels decreased by 20-30%, the reduction in fluorescence intensity was significantly more pronounced in *bud311 bud411* cells, consistent with the complete disassembly of septin structures at this stage ^29^ (note that the residual fluorescence is due to cytoplasmic mCherry-Cdc3 signal that persists in our ROI, see Materials and Methods). Recruitment of Kel1-eGFP, Sec3-eGFP and Bni1-GFP to the bud neck during cytokinesis was not affected by *BUD3* and *BUD4* double deletion (Fig. S5A-C). In stark contrast, accumulation at the bud neck of Aim44-eGFP, Nba1-eGFP, Nis1-eGFP and, to a lower extent, Spa2-eGFP, was severely impaired in *bud311 bud411* cells relative to their wild type counterpart (Fig. 7A-D). In particular, these proteins appeared at the bud neck around the same time in wild type and mutant cells, but failed to rise to high levels in the absence of Bud3 and Bud4, suggesting that membrane compartmentalization by the double septin ring may help to efficiently concentrate these proteins at the division site. This was particularly obvious for Nba1-eGFP, which was initially recruited to similar levels in wild type and *bud311 bud411* cells at the time of cytokinesis, but then quickly dissipated (Fig. 7B), presumably by diffusion along the plasma membrane.

**Figure 7.**
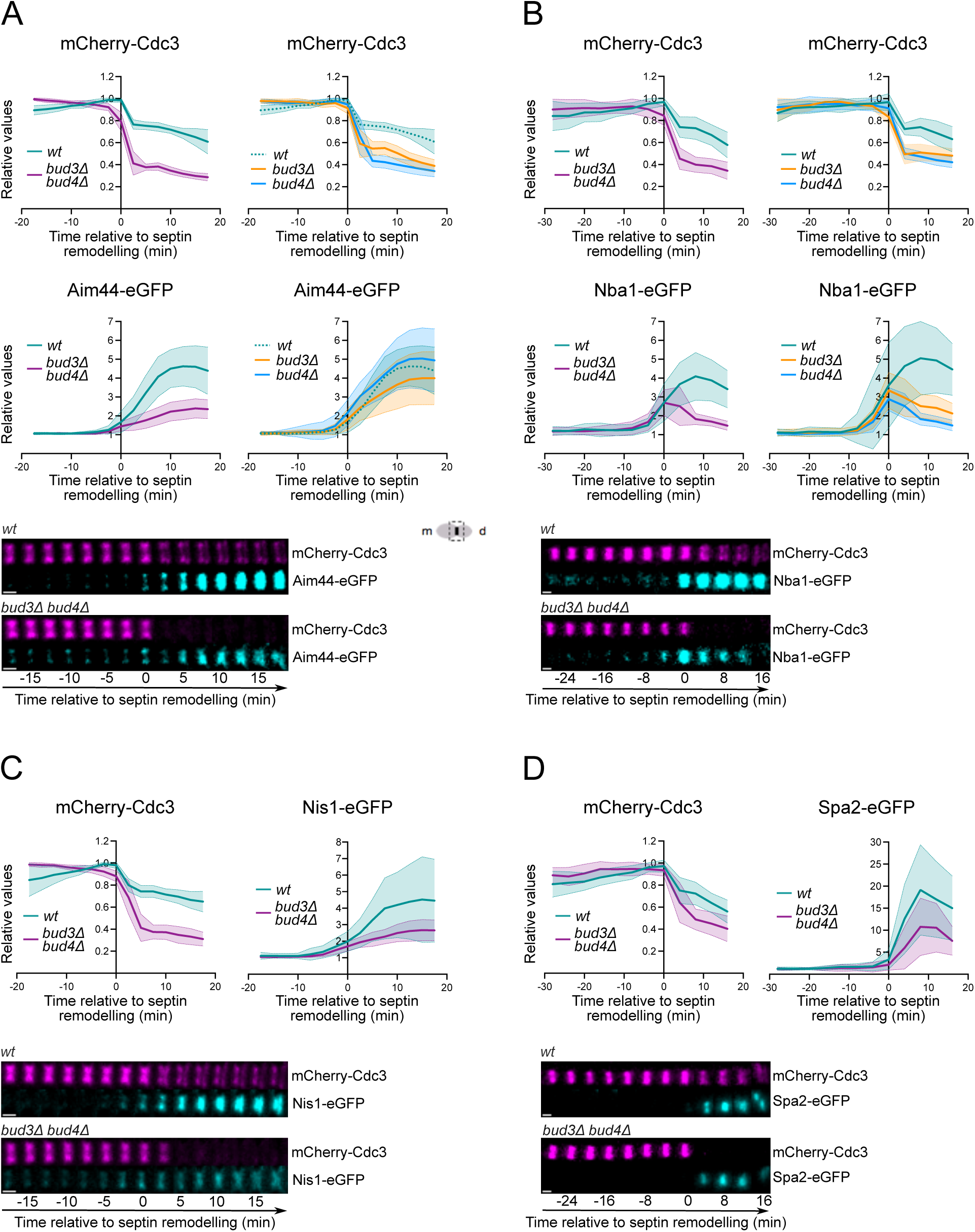
The double septin ring limits the diffusion of some bud-neck localizing proteins. **A-D:** Wild type or *bud3Δ*, *bud4Δ* and *bud3Δ bud4Δ* cells expressing mCherry-Cdc3 and the indicated eGFP-tagged protein were filmed at 30°C every 2.5 minutes (Aim44-eGFP and Nis1-eGFP) or every 4 minutes (Nba1-eGFP and Spa2-eGFP) to quantify the mean fluorescence intensity over time of mCherry-Cdc3 and the eGFP-tagged protein before and after septin remodelling as detailed in Materials and Methods. Values were averaged and plotted. Shaded curves: standard deviations. Representative frames of fluorescent signals at the bud neck are shown at the bottom. Scalebar: 1 µm.

We then studied the distribution of Aim44-eGFP and Nba1-eGFP also in the single mutants *bud311* and *bud411.* Deletion of *BUD4* had no effect on Aim44-eGFP bud neck recruitment, while deletion of *BUD3* modestly decreased its gathering at the bud neck (Fig. 7A), indicating that Bud3 and Bud4 share partially overlapping roles in constraining Aim44 at the division site. In contrast, deletion of either *BUD3* or *BUD4* severely impacted the bud neck retention of Nba1-eGFP, similar to the double deletion (Fig. 7B), suggesting that both Bud3 and Bud4 are necessary to hold Nba1 at the bud neck.

Thus, the membrane boundary generated by the double septin ring formed by Bud3 and Bud4 may selectively prevent the diffusion of some, but not all proteins that localize between the two septin rings.

## DISCUSSION

The septin hourglass to double ring transition involves a remarkable rearrangement of septin filaments that is essential for cytokinesis. However, its underlying molecular mechanism remains ill-defined.

Bud3 and Bud4 have long been known to localize in mitotic cells as a double ring at the two edges of the septin hourglass and to remain in this configuration well after septin remodelling, when they precisely colocalize with the double septin ring ^29,31,32,39,42,51,53^. These observations, together with the phenotypic characterization of *bud3* and *bud4* mutants, led to the proposal that Bud3 and Bud4 provide a spatial cue to instruct the assembly of the double septin ring at cytokinesis ^29^. Our FRAP data suggest that Bud3 are Bud4 are not required for the assembly of circumferential filaments, which may already be present in the septin hourglass in mitosis. Rather, Bud3 and Bud4 may stabilise circumferential filaments at the moment of cytokinesis to preserve them from the complete disassembly of the septin collar.

By using *in vitro* reconstitution assays, we show here that Bud3 and Bud4 can each induce the formation of septin higher order assemblies in solution. However, the two proteins have different effects on septin organisation, with Bud3 promoting bundling of septin filaments and Bud4 inducing the lateral bridging of paired septin filaments. We note that although we occasionally observe paired septin filaments with regularly spaced lateral connections in solution after long polymerisation times, the presence of Bud4 greatly stimulates their formation. Thus, while the lateral bridges are presumably made by one specific septin (most likely Cdc11), Bud4 may stabilize them or impose a septin conformation that is favourable to bridging. Interestingly, platinum-replica electron microscopy experiments previously showed that circumferential filaments in the transitional septin hourglass displayed Bud4-dependent bulging bridges approximately spaced by the length of one septin octamer ^29^, suggesting that Bud4 could indeed assist in crosslinking adjacent septin filaments. In line with this conclusion, we show that Bud4 co-localizes with septin structures that are generated *in vitro* by Bud4 alone or Bud3 and Bud4 together.

In our *in vitro* assays, combined addition of Bud3 and Bud4 leads to synergistic septin bundling and membrane interaction, suggesting that the two proteins modulate septin architecture in a highly cooperative manner. These results are consistent with the notion that Bud3 and Bud4 play distinct and complementary roles in the formation of the double septin ring *in vivo* ^29^. This conclusion is further strengthened by the evidence that Bud3 and Bud4 have different septin-binding specificities, with Bud3 interacting preferentially with Cdc10, while Bud4 was previously found to bind Cdc3, Cdc11 and Shs1 ^51^. Additionally, we find that Bud3 and Bud4 interaction with each other is mediated by septin filaments, since their association is lost upon septin depolymerization in the *cdc12-*6 temperature-sensitive mutant. Altogether, our data indicate that both Bud3 and Bud4 have the ability to bind directly septin filaments independently of one another. Consistently, Bud3 still localises at the bud neck of *bud411* cells, albeit at reduced levels, and reciprocally Bud4 localises efficiently at the bud neck of *bud311* cells ^29^ (this manuscript). The decreased association of Bud3 with the septin collar and its loss from the bud neck at the time of cytokinesis in *bud411* cells suggest that Bud3 may have high affinity for Bud4-induced septin structures, in agreement with our *in vitro* data.

How do Bud3 and Bud4 interact with septins? Previous studies defined a minimal domain of Bud4 (aa 623-774) as being involved in septin binding ^32,51^. However, our data show that a naturally occurring variant of Bud4 (*bud4-G2459fs*) that retains this domain but lacks 609 aa at the C-terminus almost abolishes Bud4 recruitment to the bud neck. Deletion of the PH domain alone (aa 1286-1447) resulted in abnormal septin remodelling, with septin disassembly on the mother side of the bud neck, and affected Bud4 localisation at the septin collar but not at the septin structures remaining after cytokinesis, in line with published results using a deletion mutant that eliminates both the anillin-homology and PH domains ^29^. Thus, the anillin-homology domain is unlikely to contribute to septin binding, while the PH domain together with the minimal domain in the middle promotes efficient Bud4 interaction with septins, probably aided by other determinants located between aa 838 and 1286 (i.e. the region missing in *bud4-G2459fs* cells). Interestingly, a recent paper highlighted the involvement of Bud3 amphipathic helix (aa 847-865) together with a domain at the C-terminus (aa 1172-1273) in the recruitment of Bud3 to septins ^43^. Thus, the combined contribution of a membrane-interacting domain and the C-terminus that was previously implicated in Bud3-Bud4 association ^32,37^ seems to be a common feature shared by Bud3 and Bud4 for septin binding.

Here we show that Bud3 and Bud4 are retained at the bud neck well after septin disassembly in a septin mutant that fails to form the septin double ring at cytokinesis, suggesting that they might interact directly with the membrane independently of septins in yeast cells. Additionally, using lipid monolayers or supported lipid bilayers combined with fluorescence microscopy or TEM/cryoEM, we show that Bud3 and Bud4 can remodel membranes *in vitro* by inducing deformations. Deletion of the amphipathic helix of Bud3 or the PH domain of Bud4 does not abolish membrane deformations, but rather decreases their radius, consistent with the idea that PH domains and amphipathic helices can modulate membrane curvature ^71–73^. In turn, this could potentially impact the curvature preference of septin filaments. Since Bud3 and Bud4 stabilise the circumferential filaments of the septin double ring, which lays on the sides of the bud neck where the plasma membrane has shallower curvature. Whether and how Bud3 and Bud4 directly interact with the plasma membrane *in vivo* and how they modulate the curvature preference of septin filaments are important questions that remain to be addressed.

Importantly, while septin remodelling or clearance from the bud neck is essential for cytokinesis^27^, the double septin ring is not. Indeed, *bud311 bud411* double mutants constrict the actomyosin ring with wild type kinetics despite complete disassembly of septin structures at the bud neck during cytokinesis ^29^. An obvious question then arises: what is the functional meaning of the septin double ring? The septin double ring was proposed to create a boundary that compartmentalizes the plasma membrane around the cleavage site ^17^. Our data show that some, but not all, proteins that are recruited to the bud neck around the time of cytokinesis need a double septin ring to concentrate efficiently at the bud neck. In particular, Aim44, Nis1, Nba1 and Spa2 seem to gather at the bud neck of *bud311 bud411* cells without reaching the peak levels observed in the wild type. In contrast, we did not observe any obvious difference in the rise of Kel1, Sec3 and Bni1 at the bud neck in *bud311 bud411* compared to wild type cells. Particularly revealing is the behaviour of Nba1, which arrives at the bud neck shortly before septin remodelling and then quickly dissipates in *bud311 bud411* cells, while it keeps rising in wild type cells. This behaviour is consistent with the proposal that septins can function as diffusion barriers ^16–18,74–78^, while it argues against a role of Bud3 and Bud4 in scaffolding the Aim44-Nis1-Nba1 complex at the bud neck. Interestingly, Nba1, Aim44 and Nis1 are all part of a protein complex that prevents rebudding at the old bud site by keeping Cdc42 activity locally low ^67,79,80^. These proteins localize at the division site and, after cytokinesis, remain at cytokinesis remnants well after septins have been disassembled to act as a spatial memory cue that prevents the reuse of an already used polarity site. In the absence of such a surveillance mechanism the replicative lifespan of budding yeast cells is considerably shorter ^67^. In the future, it will be interesting to assess if *bud311 bud411* mutant cells re-bud at the old bud site, as previously shown for *aim44Δ* cells, and whether inefficient recruitment of the Aim44-Nis1-Nba1 complex to the bud neck has any link with the known bud site selection defects of *bud*3 and *bud4* mutants ^33,33,37,53^. Interestingly, Bud4 has been proposed to act as an early landmark for the specification of future septation sites in *N. crassa* ^81^, suggesting that anillins may bear a conserved memory function for polarized processes.

We find that Spa2, but not other polarisome components, is also recruited less efficiently to the bud neck of *bud311 bud411* mutant cells relative to the wild type control, suggesting that its retention at the division site may be optimized by a septin-dependent diffusion membrane, as previously proposed ^17^. Spa2 has been involved in cell polarity, cell wall integrity and cytokinesis ^82–85^. Interestingly, it has also been implicated in bud site selection ^86^. Whether the failure to properly accumulate Spa2 at the bud neck can partially account for the abnormal budding pattern of *bud3* and *bud4* mutant cells is an intriguing possibility that will be addressed in the future.

Altogether, our data suggest that the septin diffusion barrier at the bud neck is selective towards specific set of proteins. How could selectivity be achieved? The proteins we have scrutinised cover a wide range of molecular weights (46-220 KDa) and isoelectric points (4.82-10.47), and no correlation was found with their ability to be or not constrained by the double septin ring at the bud neck, suggesting that these parameters are unlikely to underlie the selectivity of the septin diffusion barrier. Rather, we speculate that these proteins may be recruited to the bud neck through direct or indirect interactions with specific lipids at the plasma membrane, which in turn may be concentrated at the cleavage site by a septin diffusion barrier. For instance, septins have been recently shown to limit the lateral diffusion of PI(4,5)P_2_ at the plasma membrane ^87^.

A septin double ring has been found at the cleavage site of other cell types, such as fission yeast, filamentous fungi and mammalian cells ^45,46,88–90^, raising the possibility that the principles governing its assembly may be conserved. The finding that the anillin Mid2 colocalises with and stabilises the double septin ring in *S. pombe* ^45,46,88^ supports this idea.

## MATERIALS AND METHODS

### Yeast strains, growth conditions and plasmids

Most yeast strains (listed in Table S1) share the same W303 genetic background (*ade2-1, trp1-1, leu2-3,112, his3-11, 15 ura3*). Strains used for the tripartite GFP system were in the BY4741 background (*his3Δ1, leu2Δ0, met15Δ0, ura3Δ0*). Strains used for protein purifications are derived from crossing CG378 and S288c. W303 carries a frameshift mutation in the *BUD4* gene (*bud4-G2459fs*) that due to a premature stop codon results in the production of a truncated protein of 838 amino acids lacking 609 amino acids and carrying 18 non-natural amino acids at the C-terminus. This mutation disrupts septin remodelling at cytokinesis, causing septin disassembly instead ^30^. To overcome this issue and enable proper visualization of septin double rings, yeast strain genotypes were corrected to carry full-length *BUD4* whenever relevant ^27,91^.

Yeast cultures were grown at 30°C in either synthetic medium supplemented with the appropriate nutrients or YEP medium (1% yeast extract, 2% bactopeptone, 50 mg/l adenine). Raffinose was supplemented to 2%, glucose to 2% and galactose to 1%.

One-step tagging techniques ^92,93^ were used to generate strains bearing *bud3::HPH*, *BUD3-3HA, BUD4-6Gly-3Flag, BUD3-mNeonGreen, BUD4-4GFP (*full-length and Δ1286-1447*), BUD4-eGFP (*full-length and Δ1286-1447*), AIM44-eGFP, NBA1-eGFP, NIS1-eGFP, SPA2-eGFP, SEC3-6Gly-eGFP* and *BUD3-linker(3aa)-GFPβ11* ^52^. The *bud3::HPH* strains do not carry a complete deletion of the gene, but they leave intact the N-terminal GEF domain of the protein (aa 1-443) that is not involved in the septin double ring stability ^29^. The *bud4::HIS3* deletion strain was a kind gift from E. Bi and was backcrossed at least four times to W303. To generate Bni1-GFP, we integrated at the endogenous *BNI1* locus an integrative plasmid carrying the 3’ end of *BNI1* ORF fused to the coding sequence of GFP ^94^. A W303 strain carrying the *KEL1-GFP* fusion was a generous gift from D. Panigada and M. Muzi-Falconi. A W303 strain carrying *mCherry-CDC3* was a kind gift from A. Amon ^95^.

The *MBP-BUD3* and *MBP-BUD4ΔN* fusions for Bud3 and Bud4 purification from yeast cells were generated using an auto-selection expression system ^96^. Briefly, PCR cassettes encoding full-length *BUD3* or *BUD4ΔN* along with gapped pMG3 expression vector were used to transform the yeast expression host MGY853 (*MATa ura3-1 trp1-28 leu2Δ0 lys2 his7 cdc28::LEU2 pep4::LYS2 [URA3-CDC28]*) by gap-repair to generate plasmid-borne *MBP-BUD3* and *MBP-BUD411N* chimeras under the control of the *GAL1-10* promoter directly in yeast cells ^96^. Auto-selection for the expression construct was achieved by passage of the resulting transformants onto 5FOA-containing medium to select for loss of the resident *CDC28*-carrying plasmid. Proteins were induced in YEP-galactose (1% final) for 8 hr at 30 °C. Cells were harvested and proteins purified on amylose beads as previously described ^97^. To make sure that no endogenous Bud3 would copurify along with Bud4 and vice-versa, for all experiments we deleted *BUD4* in the *MBP-BUD3* strain and *BUD3* in the *MBP-BUD4ΔN* strain. Any additional tags or mutations were added to the *MBP-BUD3* and *MBP-BUD4ΔN* plasmids in yeast. Membrane-binding domain mutants of *BUD3* and *BUD4* (deletion and point mutations) were made by mutagenesis or overlap extension PCR. Specifically, *BUD3*-ΔAHX (Δaa850-858) and *BUD4*-ΔPH (Δaa1285-1424) were constructed by PCR excluding the membrane binding domains and used for end-on ligation. *BUD3-AHXm* (L854R, V857Q, F858N) was made by overlap extension PCR, while *BUD4-PHm* (R1323N, R1340N, K1341N) was made by two subsequent rounds of site-directed mutagenesis. The HALO tag was added to the C-terminus of MBP-Bud411N using PCR and Gibson assembly. The plasmids were used for bacterial transformation and finally used to retransform the yeast expression host MGY853.

For the tripartite GFP system ^52^, haploid strains carrying *BUD3-linker(3aa)-GFPβ11* and a plasmid to express *GFPβ1-9* from the *GAL1* promoter, were crossed with haploid strains carrying GFPβ10-tagged *CDC3*, *CDC10*, *CDC12*, *CDC11* or *SHS1* and expressing Cdc11-mCherry, a kind gift from J. Thorner ^51^.

### Protein expression and purification

Recombinant Cdc11-capped septin octamers were isolated as previously described ^48^. In short, *E.coli* strain BL21 (DE3) Rosetta cells were transformed with either the bicistronic plasmids pFM453 (*colE1*, *Amp^r^*, *CDC10*, *CDC11*) and pFM455 (p15A, *Kan^r^*, *MBP-CDC12*, *His6-CDC3*), or the bicistronic plasmids pFM453 and pFM873 (p15A, *Kan^r^*, *MBP-CDC12*, *His6-CDC3-yeGFP*) ^47^. Freshly transformed cells were grown in LB containing 50 µg/ml of ampicillin, 25 µg/ml of kanamycin, 34 µg/ml of chloramphenicol, 0.2% glucose, and induced at OD600=1 by adding 0.2 mM IPTG. After 20 hours at 16°C, cells were harvested by centrifugation and resuspended in Buffer A (25 mM NaHPO_4_ pH 7.8, 300 mM NaCl, 0.5 mM MgCl_2_, 5% glycerol) containing 5 mM β-mercaptoethanol, 1 mg/ml of lysozyme, and protease inhibitors (Complete EDTA-free Roche). Cell extracts were obtained by sonication and centrifugation at 30,000 *g* for 30 minutes at 4°C. The lysates were incubated with 1.5 ml amylose resin (New England Biolabs) in buffer A on a rotating wheel for 2 hours at 4°C. The slurry was washed 3 times with buffer A and loaded onto a Polyprep column (Bio-Rad). Fractions of 0.5 ml were eluted with buffer A containing 10 mM maltose and quantified using a Nanodrop. The fractions with the highest protein concentrations were combined and MBP was cleaved from the complex using thrombin at 5 U/mg overnight at 6°C. Proteins were diluted in 10 ml of buffer B (25 mM NaHPO_4_ pH 7.8, 300 mM NaCl, 0.5 mM MgCl_2_, 5% glycerol, 0.1% Triton X100, 6 mM imidazole) supplemented with 5 mM β-mercaptoethanol and incubated with 1 ml of pre-washed Ni-NTA agarose beads (Qiagen) on a rotating wheel for 2 hours at 4°C. The slurry was then washed 3 times with buffer C (25 mM _NaHPO4_ pH 7.8, 500 mM NaCl, 0.5 mM MgCl_2_, 5% glycerol, 0.15% Triton X100, 5 mM β-mercaptoethanol, 8 mM imidazole) and loaded onto a Polyprep column (Bio-Rad). Fractions of 0.5 ml were eluted with buffer B containing 200 mM imidazole and quantified using a Nanodrop. The most concentrated fractions were dialyzed against a buffer containing 20 mM Tris-Cl pH 8.2, 300 mM NaCl, 0.2 mM MgCl_2_, and 2 mM DTT. Proteins were finally concentrated using Amicon Ultra filter units (10 kDa cut-off), aliquoted and snap-frozen. The concentration of the protein was finally determined using gel electrophoresis and Coomassie staining with BSA standard curve.

To purify recombinant 6His-Bud4, *E.coli* strain BL21 (DE3) Rosetta cells were transformed with a pPROEX HTa plasmid for expression of Bud4(623-1447). Cells were grown overnight in LB containing 50 µg/ml of ampicillin, 34 µg/ml of chloramphenicol, 0.2% glucose, and induced at OD600=1 by adding 0.2 mM IPTG. After 20 hours at 16°C, cells were harvested by centrifugation and resuspended in lysis buffer (50 mM Tris-Cl pH 8, 300 mM NaCl and 0.1% Tween20) supplemented with protease inhibitors (Complete EDTA-free Roche), 10 mM imidazole and 1 mg/ml of lysozyme. Cell extracts were obtained by sonication and centrifugation at 30,000 g for 30 minutes at 4°C. The lysates were incubated with 1 ml Nickel-NTA agarose (Invitrogen) in lysis buffer containing 10 mM Imidazole on a rotating wheel for 2 hours at 4°C. The slurry was washed 3 times with lysis buffer containing 20 mM imidazole and loaded onto a Polyprep column (Bio-Rad). Fractions of 0.5 ml were eluted with buffer B containing 200 mM imidazole and quantified using a Nanodrop. The most concentrated fractions were dialyzed against a buffer containing 20 mM Tris-Cl pH 8.2, 300 mM NaCl, 0.2 mM MgCl_2_, and 2 mM DTT. Proteins were finally concentrated using Amicon Ultra filter units (50 kDa cut-off), aliquoted and snap-frozen. The concentration of the protein was finally determined using gel electrophoresis and Coomassie staining with BSA standard curve.

MBP-Bud3 and MBP-Bud4 were produced in budding yeast using an auto-selection expression system previously developed ^96^. Proteins were induced in YEP-galactose (1% final) for 8 hours at 30 °C. Cells were harvested and proteins purified on amylose beads as previously described ^97^. In short, cells were harvested by centrifugation, washed and resuspended in Lysis buffer (50 mM Tris-Cl pH 7.5, 250 mM NaCl, 0.2% Triton X100, 10% Glycerol, 5 mM EDTA) supplemented with protease inhibitors (Complete EDTA-free Roche). Glass beads were added and samples vigorously shaken in dry ice with a FastPrep (MP Biomedicals) for 5 rounds of 20 seconds interrupted by cooling for 4 minutes with dry ice. Lysates were harvested and ultra-centrifuged at 50,000 rpm for 1 hour at 4°C. The cleared lysates were incubated with 1 ml amylose resin (New England Biolabs) on a rotating wheel for 2 hours at 4°C. The slurry was washed 3 times with wash buffer (50 mM Tris-Cl pH 7.5, 250 mM NaCl and 0.2% Triton x100) and loaded onto a Polyprep column (Bio-Rad). In the case of Bud4-HALO, half of the slurry was labelled with 2 μM HaloTag® Alexa-Fluor® 660 fluorescent ligand (Promega) for 2 hours on ice, and washed once with wash buffer before loading onto a Polyprep column (Bio-Rad). Fractions of 0.5 ml were eluted with wash buffer containing 20 mM maltose and quantified using a Nanodrop. The fractions with the highest concentrations were combined and dialyzed against a buffer containing 50 mM Tris-Cl pH 7.5, 250 mM NaCl, 0.5 mM DTT. Proteins were finally concentrated using Amicon Ultra filter units (50 kDa cut-off), aliquoted and snap-frozen. The concentration of the protein was determined using gel electrophoresis and Coomassie staining with a BSA standard curve.

### Liposome preparation and Supported Lipid Bilayer preparations

Liposomes were prepared by dehydrating a lipid mixture of 87 mol% Egg-PC, 2.8 mol% Brain PI(4,5)P_2_, 10 mol% Liver-PI and 0,2 mol% TF-TMR-PI(4,5)P_2_ or 95.8 mol% DOPC, 4 mol% DGS-NTA(Ni) and 0.2 mol% Rhodamine-PE (Avanti Polar Lipids) in a vacuum oven at 60°C. Lipids were rehydrated with Citrate buffer (20 mM Citrate pH 4.6, 50 mM KCl, 0.5 mM EGTA). Liposomes were formed by vortexing and incubating at 44°C. Lipid mixtures were passed 15 times through a lipid extruder with a 100 nm pore size on a heated plate at 70°C (internal temperature 40°C) to produce small uniform-sized liposomes. A final concentration of 0.2 mM lipids was used in case of flat SLBs and 1 mM in case of SLBs on silica beads.

Coverslips were cleaned by cycles of 30-minute sonication in NaOH 1M, 100% ethanol and milliQ water before storage in 100% ethanol. On the day of the experiment, the slides were air dried and plasma cleaned in a Femto plasma surface cleaner (Diener electronic).

For flat Supported Lipid Bilayer (SLB) preparation, either 2-well Ibidi silicon culture-inserts (IBIDI) were placed on the coverslips to create a well or an imaging chamber was created by placing strips of parafilm on a microscopy slide and placing a coverslip on top, into which the SUVs were added. For fluorescence microscopy experiments, supported lipid bilayers were prepared as described previously ^98^. For the preparation of SLBs on silica beads, 20 µl SUVs were incubated together with silica beads of 1, 3, 5 μm diameter in 80 µl citrate buffer (citrate 20 mM pH 4,6, KCl 50 mM, EGTA 0.5 mM) for 30 minutes at 37°C with shaking. The beads were washed 3 times with 80 µl wash buffer (20 mM Tris pH8, 50 mM NaCl). 5 µl of beads suspension was added to a μ-Slide (8 well ibiTreat; IBIDI) in wash buffer containing 40 μM GTP, 2 mM MgCl_2_ and 1 mg/ml BSA.

### In vitro septin polymerisation assays

*S*eptin polymerisation assays were performed in solution or on an SLB with a flat support or beads. Purified septin octamers (Cdc10, Cdc11, Cdc12 and His6-Cdc3) were added together with MBP-Bud3 and/or MBP-Bud411N-HALO or membrane binding mutant variants in 20 mM Tris pH8, 2 mM MgCl_2_, 30-50 mM NaCl, 40 µM GTP and 1 mg/ml BSA, with the exception of Fig. 4 where no MgCl_2_ was added. For the experiment in Fig 1F, septins were incubated with MBP-Bud3 and/or MBP-Bud411N-HALO for 1 hour. For the experiments on SLBs, septins with or without Bud3/Bud411N were added to the SLB after a 5-minute incubation at room temperature in buffer containing 30 mM of NaCl and imaged roughly one hour after being added to the SLB (Figure 3A and 4C).

### Fluorescence microscopy and data analysis

For live cell imaging, cells were mounted on 1% agarose pads in SD medium on Fluorodishes and filmed at 30°C either with a 100X Plan TIRF Apochromat 1.49 NA oil immersion objective mounted on a Nikon Eclipse Ti microscope equipped with an EMCCD Evolve 512 camera (Photometrics) and iLAS2 module (Roper Scientific) and controlled by Metamorph (Fig. 7B, 7D, Fig. S3A-C) or with a 60X UPLXAPO 1.42 NA oil immersion objective mounted on an Olympus IX83 inverted microscope coupled to a spinning disk Yokogawa W1 and equipped with a sCMOS Fusion BT Hamamatsu camera controlled by the CellSens software (Fig. 5, 6, 7A, 7C). Z stacks of 7–12 planes were acquired with a step size of 0.4 µm and a binning of 1. Z-stacks were max-projected with ImageJ. Fluorescence intensities associated to mCherry-Cdc3 and GFP/mNeonGreen-tagged (Fig. 6, 7 and S5) were quantified with ImageJ on max-projected Z-stacks. Intensities were measured within a rectangular ROI defining the bud neck. A rolling ball radius of 50 pixels was set to subtract the background and fluorescent intensities were determined using ImageJ Analyze Particles tool after applying a threshold. For Fig. 6, the mean cytoplasmic background in a ROI of the same size as the one used for bud neck measurements was also subtracted from the mean fluorescence intensities at the bud neck. For mCherry-Cdc3, Bud3-mNeonGreen and Bud4-4GFP the highest fluorescence mean intensity reached at any time point within the time frame of interest was set at 100%, while all the other fluorescence intensities were relative to this reference value. For GFP-tagged proteins in Fig. 7 the lowest fluorescence mean intensity reached at any time point within the time frame of interest was set at 1%, while all the other fluorescence intensities were relative to this reference value. Relative intensities were then averaged after setting as time=0 the time immediately preceding septin remodelling.

For tripartite GFP imaging (Fig. 2A-D), the Cdc11-mCherry signal was used to specify a Region Of Interest (ROI) for each cell using the threshold function of Image J to quantify the tripartite GFP signal. For the analysis shown in figure 2B, the measurements on the septin collar were done in large budded cells before septin remodelling. The background GFP signal was measured inside the cell away from the septin collar and subtracted from the measurement at the septin collar. A mean intensity cut-off of 50 (dashed grey line in Fig. 2B, D) was arbitrarily set to highlight the most striking interactions.

For Fluorescence Recovery After Photobleaching (FRAP) experiments (Fig. S4), cells were imaged with a 63X 1.4 NA oil immersion objective mounted on a confocal Zeiss LSM880 Airyscan with a heated chamber at 30°C. 11 Z-stacks were acquired with a step size of 0.15 µm and later max-projected with ImageJ. Cells were arrested in metaphase by *MPS1* overexpression ^59^ to have them synchronised at the same stage of the septin hourglass. Half of the mCherry-Cdc3 septin collar was bleached with a 561 nM laser at 75% laser power for 30 iterations. In the same region, the mCherry-Cdc3 intensity was measured before and after bleaching. The background signal was measured in the cell away from the septin collar in each timepoint and subtracted from the mean mCherry intensity of the FRAP area. The mean intensity prior to bleaching was set at 100% and was used as reference to calculate relative fluorescence intensities at and after bleaching.

For *in vitro* reconstitution assays, on SLBs or in solution, fluorescent septins (Cdc11-capped octamers carrying Cdc3-GFP) were imaged with a 63X 1.4 NA oil immersion objective mounted on a confocal Zeiss LSM880 Airyscan. Z-stacks made of a variable number of planes (10-30) depending on the size of the structure were acquired with a step size of 0.15 µm and max-projected with ImageJ.

Mean intensities of the TopFluor®-TMR-PI(4,5)P2 signal in max-projected images of SLBs (Fig. S2B) were measured in ImageJ.

Septin filament length and curvature in Fig. 3A-C were analysed using the SOAX software ^99^. Curvature was analysed with 15 coarse graining snake points.

For fluorescence microscopy of septin filaments in solution (Fig.1F), 40 nM of septins were incubated together with 20 nM of Bud3 and or Bud4 for 1 hour at 4°C. Samples were dropped on a plasma cleaned coverslip, covered with a glass slide and imaged.

Membrane deformations were measured using ImageJ. In short, membrane deformations visualized by Rhodamine-PE were selected using a threshold and the particles analysed with the ImageJ plugin Filament Morphology Tool to measure the radius of the largest inscribed circle of the membrane deformations and protrusions.

### Transmission electron microscopy (TEM)

TEM with septins was performed as previously described ^6,54^. Specifically, septins were diluted to 70 nM or 90 nM (90 nM in Fig. 1B and 70 nM in Fig. 1D) in 20 µl of 50 mM Tris-HCl pH 8, 50 mM KCl, 2 mM MgCl_2_ and 40 µM GDP in the absence or presence of 70 nM or 90 nM Bud3 or Bud4, respectively, before being deposited in a well of a Teflon block. In case of lipid monolayers (Figure S2C), a drop (∼0.5 µL) of lipid solution in chloroform (87 mol% egg-PC, 3 mol% brain PI(4,5)P_2_ and 10 mol% liver-PI; Avanti lipids) was applied to the surface of the septin solution. The Teflon block was placed in a humid chamber and equilibrated overnight at 4°C. To capture the adsorbed proteins with the lipid monolayer, a grid was placed gently on the surface to have the sample stick to it. Samples were collected on 100 mesh Formvar carbon-coated grids (Delta Microscopies FCF100-Cu) and stained with 2% Uranyl Acetate in water for 1 min. Grids were imaged with a Tecnai F20 transmission electron microscope at 120KV and equipped with a Veleta camera.

For Fig. 1E, septins were thawed and diluted to a final concentration of 180 nM in 20 mM Tris-Cl pH8.2, 2 mM MgCl_2_, 40 μM GTP and 30 mM NaCl on ice. Septin octamers were incubated for 1 hour in ice in the presence of MBP-Bud3 or MBP-Bud411N (90 nM) or the two proteins together (45 nM each) before negative staining. Samples were absorbed on glow-discharged Formvar carbon-coated grids grids (Delta Microscopies FCF100-Cu) and stained with 1% uranyl acetate in water for 1 minute. Data were collected using a JEOL 1400 Flash transmission electron microscope at 120KV and equipped with a One View camera (Gatan Inc).

### Cryo-electron microscopy

A lipid mixture dissolved in chloroform (57 mol % EggPC, 10 mol % DOPE, 15 mol % Cholesterol, 10 mol % DOPS, 8 mol % PI(4,5)P2) was quickly dried using a flow of argon. The obtained lipid film was fully dried under vacuum for 30 min. Lipids were then resuspended in 20mM Tris-HCl pH 8, NaCl 50 mM, MgCl_2_ 2mM, after vortexing for 10 seconds. The obtained LUVs suspension was diluted to a final lipid concentration of 0.1 g/L lipid. Septins and MBP-Bud3 or MBP-Bud411N proteins were used at a final concentration of 25 or 125 nM for septins and varying concentrations of Bud proteins (from 20 to 200 nM). The mixtures were incubated for 1 hour at room temperature. 4 µL of solution was deposited on a glow discharged lacey electron microscopy grid (Ted Pella, USA). Most of the solution was blotted away from the grid to leave a thin film (<100 nm) of aqueous solution. Blotting was carried out on the opposite side from the liquid drop and plunge-frozen in liquid ethane at −181 °C using an automated freeze plunging apparatus (EMGP, Leica, Germany). Samples were kept in liquid nitrogen and imaged using a Tecnai G2 microscope (FEI, Eindhoven, Netherlands) operated at 200 kV and equipped with a 4kx4k CMOS camera (F416, TVIPS) or using a Glacios 200 kV FEG microscope (Thermofisher) equipped with a Falcon IVi camera (Thermofisher).

### Immunoprecipitations and western blot analysis

For immunoprecipitations of Bud4-6gly-3FLAG, cells expressing Bud3-HA with or without Bud4-6gly-3FLAG and with or without the temperature sensitive *cdc12-6* mutation ^16,17^ were grown either asynchronously or arrested in mitosis by nocodazole treatment (15 μg/ml). Cultures were then incubated at 37°C for 30 minutes to depolymerize septins at the bud neck in the *cdc12-6* strain. Cell pellets from 50 ml yeast cultures were resuspended in TBSN buffer (25 mM Tris-Cl pH 7.4, NaCl 100 mM, 2 mM EDTA, 0.1% NP40) supplemented with protein inhibitors (Complete, Roche), 2 mM sodium orthovanadate, 60 mM β-glycerophosphate and 1 mM DTT). Cells were lysed with glass beads by vigorous shaking with a vortex at 4°C with 14 cycles of 30 seconds breakage followed by 30 seconds on ice. Total extracts were cleared by spinning at 20000 *g* at 4°C and quantified by NanoDrop. A sample of 20 µl was withdrawn from the 250 µl total sample and denatured in 10 µl 3x SDS sample buffer at 100°C to serve as an input control. The remaining IP mix was incubated for 1h at 4°C on a nutator with 25 μl of protein G dynabeads (Invitrogen) preadsorbed for 90 min with 1 μl of anti-Flag M2 antibody and washed with TBSN to eliminate the excess of antibody. Magnetic beads were washed six times with TBSN buffer and bound proteins eluted with an excess of 3XFlag peptide (0.5 mg/ml in 50 mM Tris-Cl pH 8.3, 1mM EDTA, 0.1% SDS) upon incubation at R.T. for 25 min with vigorous shaking. 12.5 μl of 3X SDS sample buffer (240 mM Tris–Cl pH 6.8, 6% SDS, 30% glycerol, 2.28 M β-mercaptoethanol, 0.06% bromophenol blue) were added to the eluted proteins, followed by denaturation at 99°C for 3 min and loading on SDS–PAGE. Proteins were wet-transferred to Protran membranes (Schleicher and Schuell) overnight at 0.2 A and treated for western blotting.

For western blotting, proteins were transferred overnight at 0.2 A onto Protran membranes (Schleicher and Schuell). Specific proteins were detected using monoclonal anti-HA 12CA5 (AgroBio custom made, 1:5000), monoclonal anti-Flag M2 (F3165 Sigma, 1:5000), polyclonal anti-Cdc11 (sc-7170 Santa Cruz; 1:2000) and monoclonal anti-MBP (E8032S, New England Biolabs, 1:10000). Antibodies were diluted in 5% low-fat milk (Regilait) dissolved in TBST buffer (25 mM Tris, 137 mM NaCl, 2.68 mM KCl, 0.1% Tween-20). Secondary antibodies were obtained from GE Healthcare, and proteins were detected using a home-made luminol/p-coumaric acid enhanced chemiluminescence system.

### Statistical analysis

Statistical significance was assessed using a student’s *t-*test as indicated in the figure legends. For student’s *t-*tests, data distribution was checked visually for normal distribution of the datapoints. For Fig. 3B-C, statistical significance was assessed with Mann Witney test.

## ACKNOWLEDGMENTS

We are grateful to A. Amon, E. Bi, M. Farkasovsky, M. Muzi-Falconi, D. Panigada, M. Segal, J. Thorner for sharing yeast strains and plasmids; to the joint IGMM-CRBM “Yeast Media and Technologies” service for providing us with ready-to-use media; to members of the Piatti and Liakopoulos teams for fruitful discussions. This work has been supported by a grant of the Agence Nationale pour la Recherche (SEPTORG ANR-18-CE13-0015-01) to S.P. and L.P. and Agence Nationale pour la Recherche (CONSTRICT ANR-24-CE13-2422) to S.P. I.A. was funded by a fellowship of the Labex EpiGenMed. We acknowledge the imaging facility MRI, member of the national infrastructure France-BioImaging (https://ror.org/01y7vt929) supported by the French National Research Agency (ANR-24-INBS-0005 FBI BIOGEN).

