## supplemental figures for "Architecture and function of Bud3 and Bud4-induced septin structures"

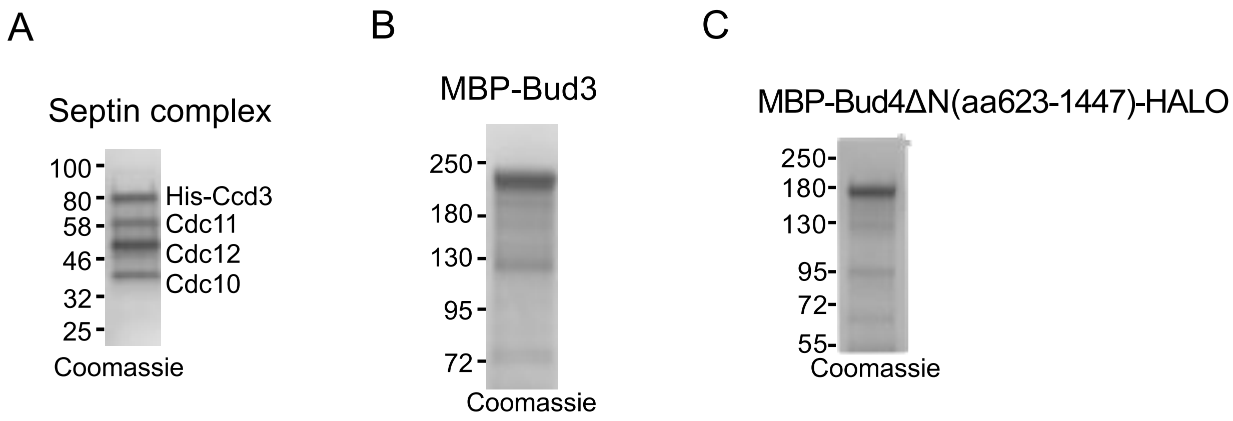


**Figure S1. Purified proteins used in this study. A:** Recombinant Cdc11-capped septin octamers were run on SDS-page and stained with Coomassie. **B:** MBP-Bud3 purified from *S. cerevisiae* and stained with Coomassie. **C:** MBP-Bud4ΔN-HALO purified from *S. cerevisiae* and stained with Coomassie.

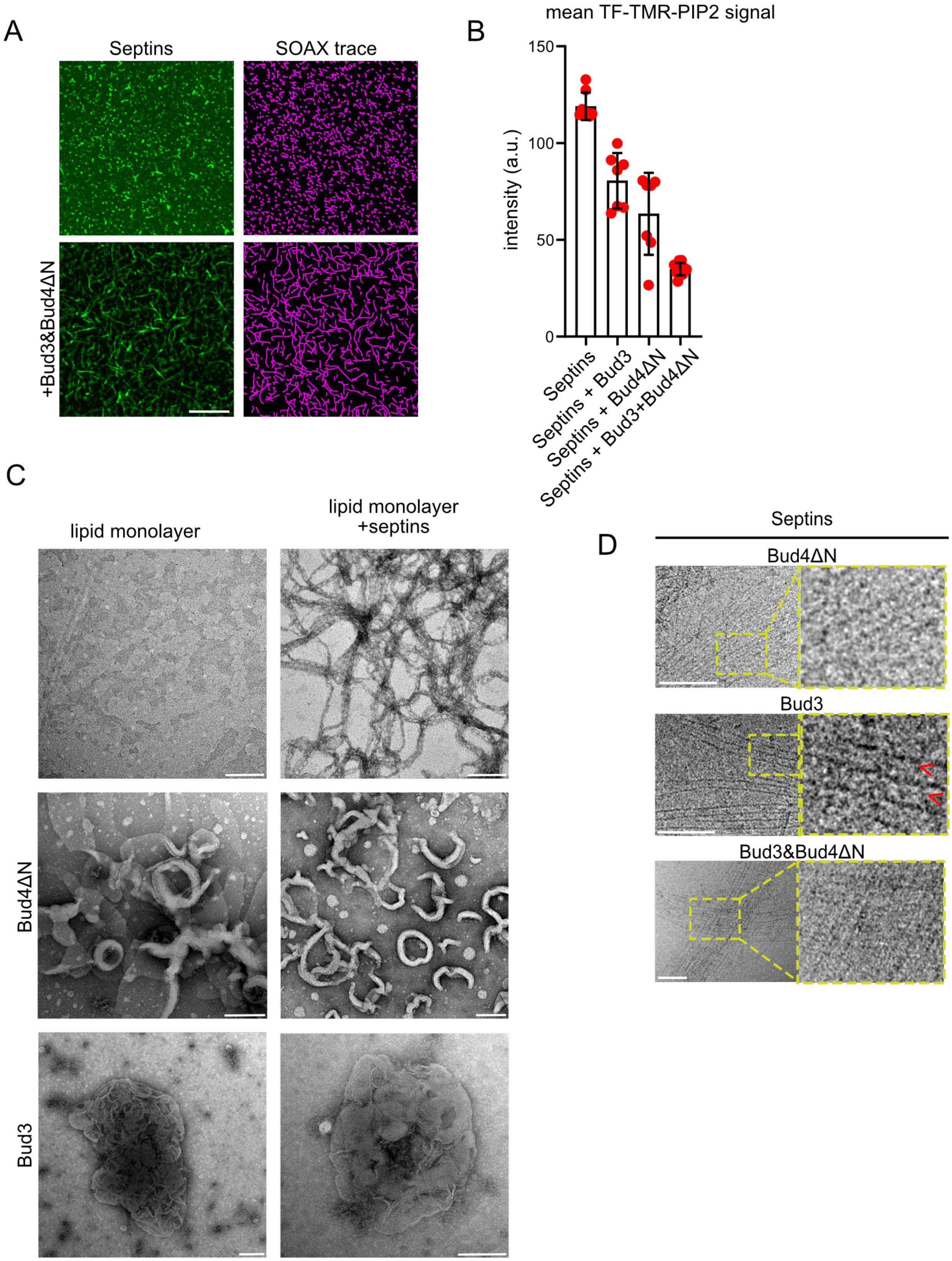


**Figure S2. Septin organization on lipid monolayers in the presence of Bud3 and/or Bud4ΔN. A:** Representative images and corresponding SOAX traces of septin structures on the SLB from the experiment in Fig. 3A-C. **B:**

Quantification of mean signals of TF-TMR-PIP2 at the SLB from the experiment in Fig. 3A-C. **C:** Representative TEM images of lipid monolayers upon addition of 70 nM Cdc11-capped septin octamers and /or equimolar Bud3 or Bud4ΔN. Scalebar: 200 nm. Note that in the presence of Bud3 and Bud4ΔN septins do not form filaments on the lipid monolayer but deformations of the monolayer become apparent. **D:** CryoEM images from the experiment in figure 3. Higher magnification images of septin bundles that are not in proximity to liposomes. Yellow boxes indicate zoom areas. Scale bars: 100 nm.

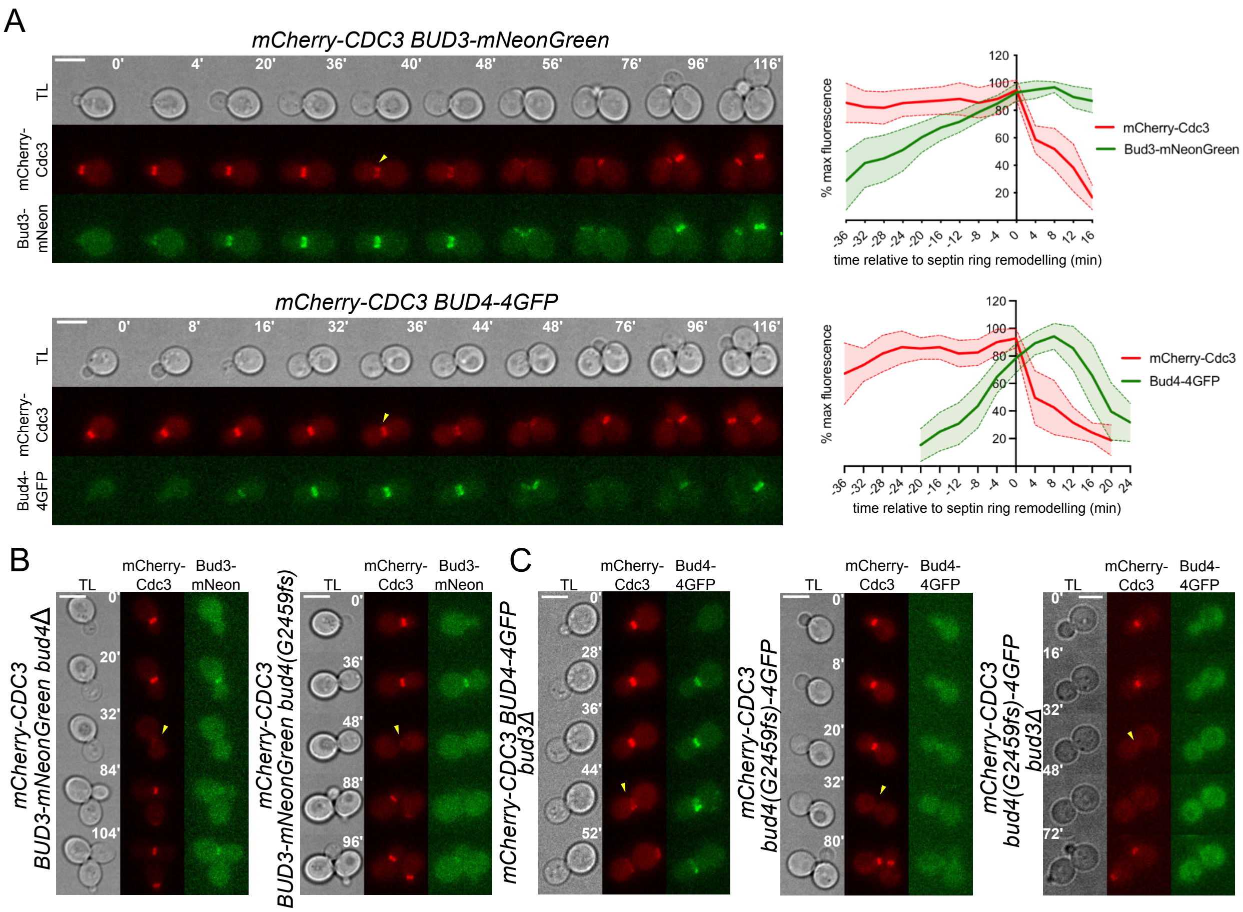


**Figure S3. Bud3 and Bud4 localisation at the septin collar is partially interdependent. A-C:** Cells expressing mCherry-Cdc3 and the indicated Bud3 or Bud4 fluorescent proteins were imaged every 4 minutes at 30°C. Fluorescent signals of mCherry-Cdc3 and Bud3-mNeonGreen or Bud4-4GFP were quantified in wild type cells. Fluorescence intensities were converted to percentages relative to the highest value for each channel and each cell, averaged and plotted relative to the time immediately preceding septin remodelling (t=0; n=20 for Bud3-mNeonGreen and n=17 for Bud4-4GFP). Shaded curves indicate standard deviations. Yellow arrowheads indicate septin remodelling. Scale bars: 5 μm.

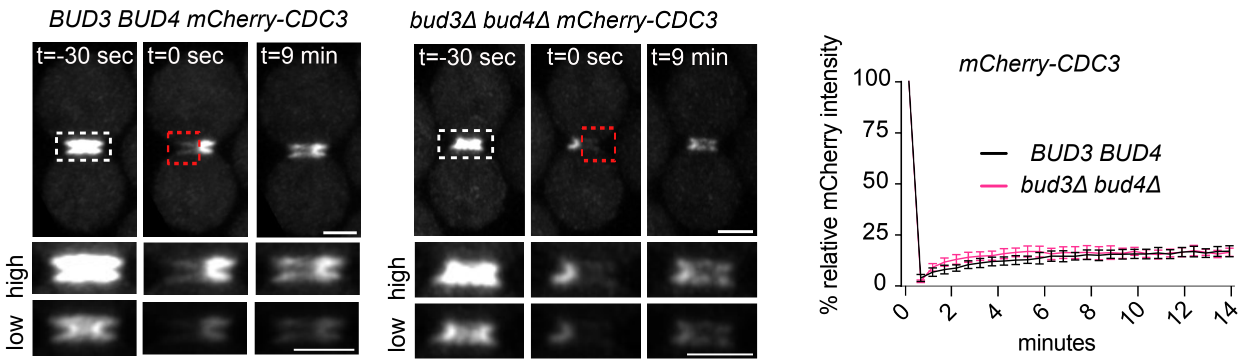


**Figure S4. A dynamic septin double ring is present at the septin collar in mitosis independently of Bud3 and Bud4.** Wild type and *bud3Δ bud4Δ* double mutant cells expressing mCherry-Cdc3 were arrested in mitosis by *MPS1* overexpression. Half of the mCherry-Cdc3 signal on the septin collar was bleached and then fluorescence recovery measured over time (graph on the right). n=11 for both conditions. Zoom areas are indicated by a white box and bleached areas by a red box. Scale bars: 2 µm. Images at the bottom row have adjusted brightness to visualize the septin collar prior to FRAP.

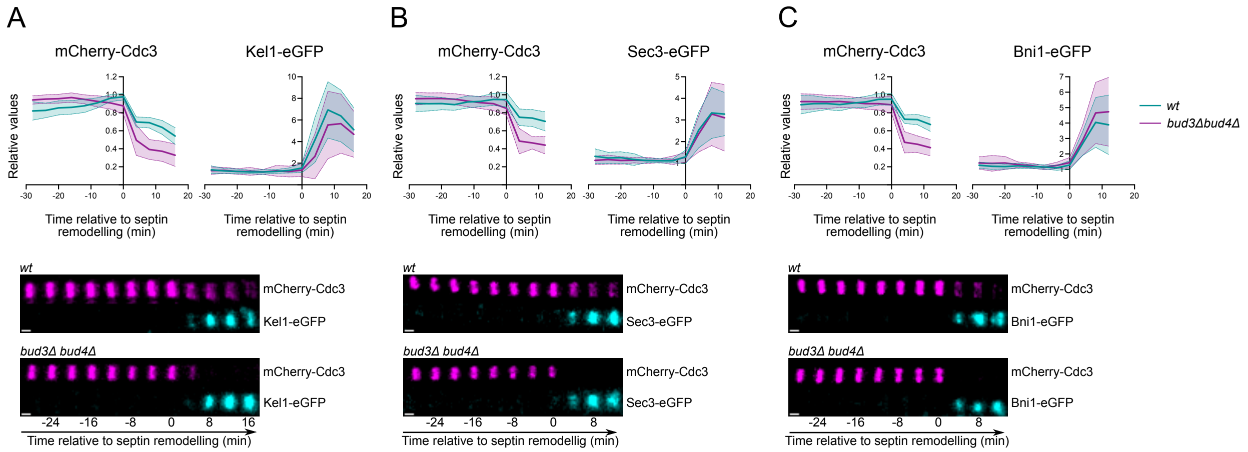


**Figure S5. The double septin ring does not concentrate all polarized proteins at the bud neck during cytokinesis. A-C:** Wild type or *bud3Δ bud4Δ* cells expressing mCherry-Cdc3 and the indicated GFP-tagged protein were filmed to quantify the fluorescence intensity over time of mCherry-Cdc3 and the GFP-tagged protein before and after septin remodelling. Representative frames of fluorescent signals at the bud neck are shown at the bottom. Scalebar: 1 µm.

**Table S1. List of *S. cerevisiae* strains used in this study**

| Name | Relevant genotype |
|---|---|
| ySP15353 | *MATa, bud4(G2459fs)::LEU2::BUD4, BUD3-mNEONgreen::NAT, cdc3::mCherry-CDC3::URA3* |
| ySP15361 | *MATa, bud4(G2459fs), BUD3-mNEONgreen::NAT, cdc3::mCherry-CDC3::URA3* |
| ySP15393 | *MATalpha, bud4(G2459fs)::LEU2::BUD4-4GFP::TRP1, cdc3::mCherry-CDC3::URA3* |
| ySP15395 | *MATalpha, bud4(G2459fs)-4GFP::TRP1, cdc3::mCherry-CDC3::URA3* |
| ySP15398 | *MATa, BUD3-3HA::K.l.URA3, bud4(G2459fs)::LEU2:: BUD4* |
| ySP15404 | *MATa, BUD3-mNeonGreen::NAT, cdc3::mCherry-CDC3::URA3, bud4::HIS3* |
| ySP15495 | *MATa, BUD3-3HA::K.l.URA3, bud4(G2459fs)::LEU2::BUD4-6Gly-3FLAG::KanMX* |
| ySP15538 | *MATa, bud4(G2459fs)-4GFP::TRP1, cdc3::mCherry-CDC3::URA3, bud3::HPH* |
| ySP16086 | *MATalpha, cdc28::LEU2, pep4::LYS2, [pHIS3-CDC28, MBP-BUD3]* |
| ySP16724 | *MATalpha, cdc28::LEU2, pep4::LYS2, [pHIS3-CDC28, MBP-BUD4 Ct-HALO], bud3::HPH* |
| ySP16746 | *MATalpha, cdc28::LEU2, pep4::LYS2, [pHIS3-CDC28, MBP-BUD3], bud4::HPHM* |
| ySP16747 | *MATalpha, cdc28::LEU2, pep4::LYS2, [pHIS3-CDC28, MBP-BUD3-AHm], bud4::HPHM* |
| ySP16795 | *MATa/MATalpha, BUD3-linker(5)-beta11GFP::HPH; cdc10::GFPbeta10-linker(5)-CDC10::ADH1(t)::HPH, cdc11::CDC11-mCherry::SpHIS5* |
| ySP16797 | *MATa/MATalpha, BUD3-linker(5)-beta11GFP::HPH, cdc3::GFPbeta10-linker(5)-CDC3::ADH1(t)::HPH, cdc10::CDC10-mCherry::SpHIS5* |
| ySP16799 | *MATa/MATalpha, BUD3-linker(5)-beta11GFP::HPH, cdc12::GFPbeta10-linker(5)-CDC12::ADH1(t)::HPH, cdc10::CDC10-mCherry::SpHIS5* |
| ySP16801 | *MATa/MATalpha, BUD3-linker(5)-beta11GFP::HPH, cdc11::GFPbeta10-linker(5)-CDC11::ADH1(t)::HPH, cdc10::CDC10-mCherry::SpHIS5* |
| ySP16803 | *MATa/MATalpha, BUD3-linker(5)-beta11GFP::HPH, shs1::GFPbeta10-linker(5)-SHS1::ADH1(t)::HPH, cdc10::CDC10-mCherry::SpHIS5* |
| ySP16918 | *MATalpha, cdc28::LEU2, pep4::LYS2, [pHIS3-CDC28, MBP-BUD4-ΔPH-HALO], bud3::HPH* |
| ySP16979 | *MATalpha, cdc28::LEU2, pep4::LYS2, [pHIS3-CDC28, MBP-BUD4-PHm-HALO], bud3::HPH* |
| ySP16998 | *MATalpha, cdc28::LEU2, pep4::LYS2, [pHIS3-CDC28, MBP-BUD3-ΔAH], bud4::HPHM* |
| ySP17019 | *MATa, cdc3::mCherry-CDC3::URA3, bud3::HPH, bud4::HIS3* |
| ySP17146 | *MATa, BNI1-GFP::URA3, cdc3::mCherry-CDC3::URA3, bud3::HPH, bud4::HIS3* |
| ySP17150 | *MATalpha, KEL1-GFP::KanMX, cdc3::mCherry-CDC3::URA3, bud3::HPH, bud4::HIS3* |
| ySP17174 | *MATalpha, bud4(G2459fs)::LEU2::BUD4, cdc3::mCherry-CDC3::URA3, SPA2-eGFP::KanMX* |
| ySP17176 | *MATa, bud4(G2459fs)::LEU2::BUD4, cdc3::mCherry-CDC3::URA3, KEL1-GFP::KanMX* |
| ySP17182 | *MATa, cdc3::mCherry-CDC3::URA3, bud3::HPH, bud4::HIS3, SEC3-6Gly-eGFP::KanMX* |

| | |
|---|---|
| ySP17190 | *MATalpha, bud4(G2459fs)::LEU2::BUD4, cdc3::mCherry-CDC3::URA3, BNI1-GFP::URA3* |
| ySP17192 | *MATalpha, cdc3::mCherry-CDC3::URA3, bud3::HPH, bud4::HIS3, AIM44-eGFP::KanMX* |
| ySP17193 | *MATa, bud4(G2459fs)::LEU2::BUD4, cdc3::mCherry-CDC3::URA3, AIM44-eGFP::KanMX* |
| ySP17195 | *MATa, bud4(G2459fs)::LEU2::BUD4, bud3::HPH, cdc3::mCherry-CDC3::URA3, AIM44-eGFP::KanMX* |
| ySP17197 | *MATalpha, cdc3::mCherry-CDC3::URA3, bud4::HIS3, AIM44-eGFP::KanMX* |
| ySP17200 | *MATalpha, bud4(G2459fs)::LEU2:: BUD4, cdc3::mCherry-CDC3::URA3, SEC3-6Gly-eGFP::KanMX* |
| ySP17212 | *MATa, bud4(G2459fs)::LEU2::BUD4, cdc3::mCherry-CDC3::URA3, NIS1-eGFP::KanMX* |
| ySP17215 | *MATalpha, cdc3::mCherry-CDC3::URA3, bud3::HPH, bud4::HIS3, NIS1-eGFP::KanMX* |
| ySP17220 | *MATa, bud4(G2459fs)::LEU2::BUD4, cdc3::mCherry-CDC3::URA3, NBA1-eGFP::KanMX* |
| ySP17223 | *MATalpha, cdc3::mCherry-CDC3::URA3, bud3::HPH, bud4::HIS3, NBA1-eGFP::KanMX* |
| ySP17225 | *MATalpha, cdc3::mCherry-CDC3::URA3, bud3::HPH, NBA1-eGFP::KanMX* |
| ySP17227 | *MATa, cdc3::mCherry-CDC3::URA3, bud4::HIS3, NBA1-yeGFP::KanMX* |
| ySP17235 | *MATa, cdc3::mCherry-CDC3::URA3, bud3::HPH, bud4::HIS3, SPA2-eGFP::KanMx* |
| ySP17314 | *MATalpha*, *bud4(G2459fs)::LEU2::BUD4-6Gly-3FLAG::KanMX, BUD3-3HA::KlURA3, cdc12-6* |
| ySP17555 | *MATalpha, bud4(G2459fs)::LEU2::BUD4ΔPH-eGFP::KanMX, cdc3::mCherry-CDC3::URA3* |
| ySP17558 | *MATalpha, bud4(G2459fs)::LEU2::BUD4-eGFP::KanMX, cdc3::mCherry-CDC3::URA3* |
| ySP18019 | *MATa, bud4(G2459fs)::LEU2::BUD4, BUD3-mNEONgreen::NAT, cdc3::mCherry-CDC3::URA3, cdc11::TRP1, ura3::URA3::CDC11ΔPB* |
| ySP18375 | *MATa, bud4(G2459fs)::LEU2::BUD4-4GFP::TRP1, cdc3::mCherry-CDC3::URA3, cdc11::TRP1, ura3::URA3::CDC11ΔPB* |
